# Engineering multivalent Fc display for FcγR blockade

**DOI:** 10.1101/2024.01.20.576357

**Authors:** Ekaterina Petrova, Georges Kiriako, Johan Rebetz, Karl Johansson, Stefan Wennmalm, Niels E.J. Meijer, B. Martin Hällberg, Ingemar André, Elena Ambrosetti, John W. Semple, Ana I. Teixeira

**Affiliations:** Department of Physiology and Pharmacology, Karolinska Institutet, Stockholm, Sweden; Division of Hematology and Transfusion Medicine, Lund University, Lund, Sweden; Department of Applied Physics, SciLifeLab, Royal Institute of Technology, Stockholm, Sweden; Department of Biochemistry and Structural Biology, Lund University, Lund, Sweden; Department of Cell and Molecular Biology, Karolinska Institutet, Stockholm, Sweden; Center for Life Nano- and Neuro-Science, Istituto Italiano di Tecnologia, Rome, Italy; Clinical Immunology and Transfusion Medicine, Office of Medical Services, Region Skåne, Lund, Sweden; Departments of Pharmacology, Medicine and Laboratory Medicine and Pathobiology, University of Toronto, Toronto, Canada

## Abstract

Autoimmune diseases, driven by Fcγ receptor (FcγR) activation through autoantibody immune complexes (IC), present a complex therapeutic challenge of achieving pharmacological blockade of FcγR without triggering receptor activation. The assembly of ICs into polydisperse, higher-order structures is required for FcγR activation. However, engineered multimeric, monodisperse Fc assemblies have been reported to prevent FcγR activation, suggesting that Fc spatial organization determines FcγR activation. In this study, we engineered a functional single-chain Fc domain protein (scFc) for unidirectional, multivalent presentation by virus-like particles (VLPs), used as a display platform. We found that the multivalent display of scFc on the VLPs elicited distinct cellular responses compared with monovalent scFc, highlighting the importance of the structural context of scFc on its function. scFc-VLPs had minimal impact on the nanoscale spatial organization of FcγR at the cell membrane and caused limited receptor activation and internalization. In contrast, the monovalent scFc acted as an FcγR agonist, inducing receptor clustering, activation, and internalization. Increasing scFc valency in scFc-VLPs was associated with increased binding to monocytes, reaching a plateau at high valencies. Notably, the ability of scFc-VLPs to block IC-mediated phagocytosis *in vitro* increased with scFc valency. In a murine model of passive immune thrombocytopenia (ITP), a high valency scFc-VLP variant with a desirable immunogenicity profile induced attenuation of thrombocytopenia. Here we show that multivalent presentation of an engineered scFc on a display platform can be tailored to promote suppression of IC-mediated phagocytosis while preventing FcγR activation. This work introduces a new paradigm that can contribute to the development of therapies for autoimmune diseases.

## Main

Protein engineering of antibody crystallizable fragment (Fc) has been instrumental in advancing antibody therapeutics. The Fc domain regulates antibody effector functions by interacting with immune cell receptors such as Fcγ receptors (FcγR) and complement system components. Mutations within the Fc domain have been designed to selectively enhance or suppress antibody effector functions, tailoring them to specific therapeutic applications.^1^ Recent advances in the field have enabled the transfer of functions previously associated with the intact IgG molecules to the Fc domain alone. Recombinant Fc-multimers,^2–6^ comprising interconnected Fc domains, have shown promise in inhibiting autoantibody-driven inflammatory reactions, characteristic of various autoimmune diseases. These multimers present an alternative to the current standard of FcγR modulation, intravenous immunoglobulin (IVIG) therapy, which is employed for treating a diverse range of diseases, including ITP,^7,8^ rheumatoid arthritis,^9^ and Kawasaki disease.^10^ However, knowledge gaps in the roles of Fc valency and spatial organization in FcγR activation, hinder further developments in the therapeutic potential and safety profile of multivalent Fc variants.

The FcγR family comprises several members, each characterized by unique expression patterns and functional properties. Notably, except for FcγRI, these receptors exhibit low-affinity interactions with monovalent IgG. Consequently, multivalent presentation of IgG becomes essential to enhance the binding affinity of the inherently weak interactions with FcγRs. The overall strength and duration of these interactions play a pivotal role in modulating the spatial reorganizationof FcγR receptors and of mediators of FcγR signaling pathways. This, in turn, can significantly impact FcγR activation levels, potentially exceeding the activation thresholds required to trigger a diverse array of immune functions, including antibody-dependent cellular phagocytosis (ADCP), antibody-dependent cellular cytotoxicity (ADCC), and platelet activation, in the case of antibody-mediated platelet diseases. Achieving the necessary affinity or avidity to engage FcγRs without inducing clustering and subsequent cellular activation poses a critical challenge, particularly in the context of preventing the binding of autoantibody-opsonized complexes. This underscores the need for investigating the roles of multivalency in Fc biology, as it holds the key to advancing the development of therapies for auto-immune diseases.

Here, by harnessing the capabilities of virus-like particles (VLPs), we have established a robust display platform. This platform facilitated the controlled presentation of engineered single-chain Fc molecules (scFc) in varying valencies. Using this approach, we developed multivalent scFc variants that could bind monocytes without inducing FcγR reorganization at the plasma membrane or receptor activation, and ultimately suppressing IC-mediated phagocytosis *in vitro* and in an animal model of passive ITP. Importantly, our findings provide a foundation for the rational design and development of novel autoimmune therapeutics.

## Results

### Engineered single-chain Fc (scFc) with Fc*γ*R binding capacity

We designed a monovalent, versatile Fc molecule tailored for site-specific conjugation to display platforms, allowing multivalent presentation. Established methods for producing monovalent Fc constructs require complex genetic and cellular engineering strategies, such as knobs-into-holes technology.^11,12^ However, these approaches often result in heterogenous products with suboptimal yield and impurities.^13^ To address the production challenges associated with the homodimeric nature of the native Fc domain, we adopted a genetic fusion approach.^1^ We redesigned the hIgG1 Fc by introducing a flexible linker between the two Fc chains to create a single chain Fc (scFc). Importantly, we retained the hinge region preceding the CH2 domain, as it plays an important role in IgG binding to Fc*γ*Rs.^14,15^ To facilitate the integration of functional modules in the scFc protein without chemical alterations, we genetically fused a SNAPtag at the C-terminus and a SpyTag at the N-terminus **(Fig 1a)**. These tags covalently bind to benzylguanine (BG) and SpyCatcher modified moieties, respectively.

**Fig. 1:**
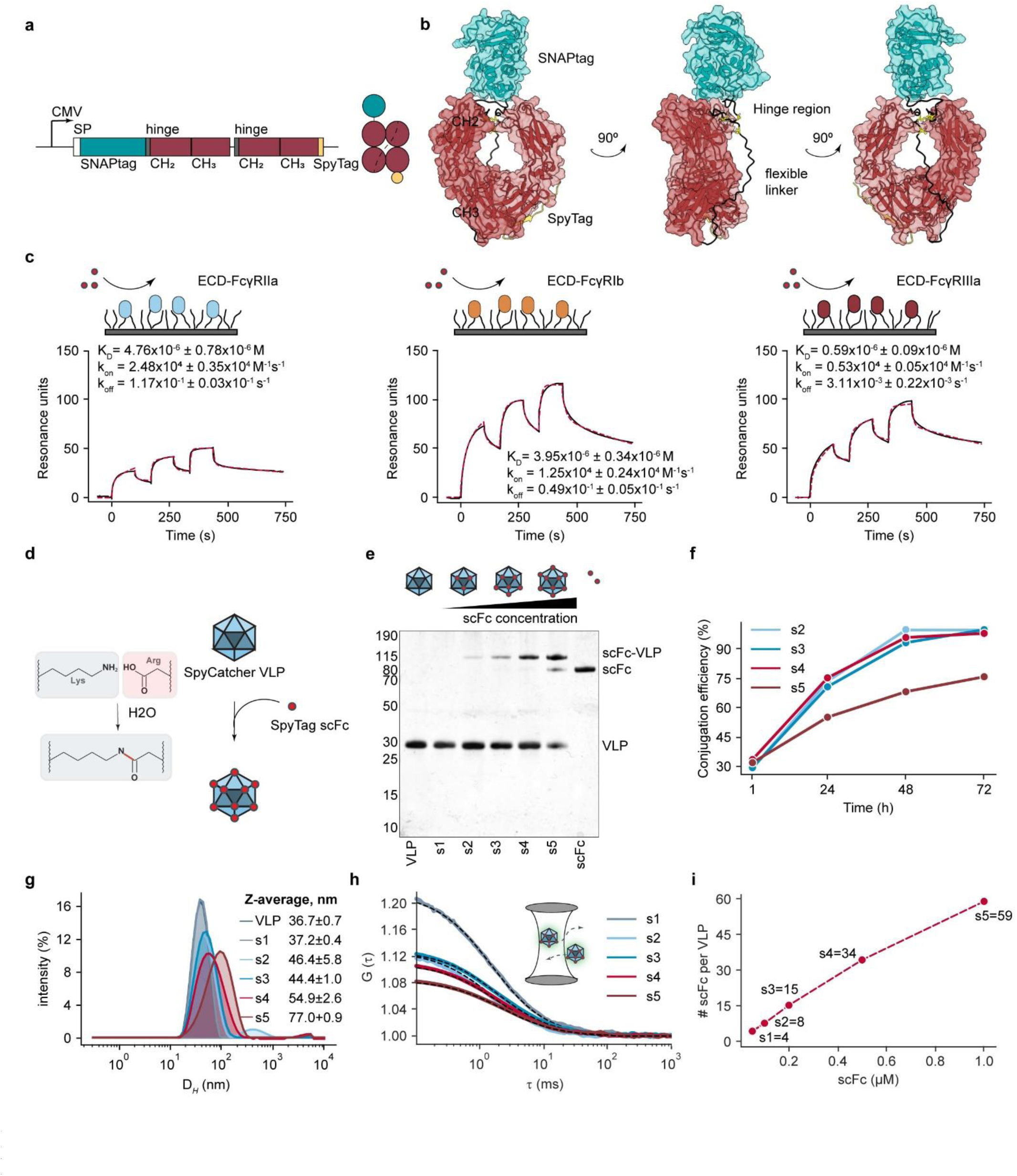
Construction and characterization of the scFc-VLP platform. **a,** Domain organization of the scFc. Different domains and regulatory elements are colored and highlighted. Two hIgG1 Fc regions, each composed of the hinge region, the CH2, and the CH3 domains, were genetically fused via a flexible linker. SNAPtag and SpyTag were added to the N- and C-terminus, respectively. Protein expression in mammalian cells was driven by a cytomegalovirus promoter (CMV), and protein secretion was induced by an optimized signal peptide (SP). **b,** Cartoon representation of the predicted scFc model. The flexible linker connecting the SNAPtag (in cyan) and the Fc region (in red) as well as the Fc interdomain flexible linker are depicted in black. The Cys residues located at the reconstructed hinge region are shown in yellow. **c,** Schematics of the SPR experimental designs and corresponding sensorgrams. ECD-FcγRIIa, ECD-FcγRIIb, or ECD-FcγRIIIa were covalently attached to an activated dextran matrix, and scFc binding was assessed using single-cycle kinetic analysis, where the analyte was injected in three increasing concentrations (0.15, 0.44, and 1.3 µM). Black lines represent the measured curves and red lines represent the bivalent analyte binding model curve fits. One representative experiment of n=2 independent repeats is shown. The dissociation constant (K_D_) and kinetic rate constants are shown as mean ± std. **d,** Schematic diagram of the scFc-VLP platform, consisting of SpyCatcher-VLP and SpyTag-scFc. Lysine and arginine residues present in SpyCatcher and SpyTag, respectively, spontaneously react to form covalent isopeptide bonds. **e,** Silver stained SDS–PAGE of VLP, scFc-VLP variants (s1, s2, s3, s4, and s5), and scFc. **f,** Conjugation efficiencies of scFc-VLP variants (s2, s3, s4, and s5) at 1 h, 24 h, 48 h, and 72 h reaction times, estimated using SDS-PAGE densitometry (n=1). **g,** DLS characterization of VLP and scFc-VLPs (n=3, mean ± std). D*_H_*, hydrodynamic diameter. **h,** Autocorrelation curves derived from FCS measurements on ATTO 488-labeled scFc-VLPs. One component fits are shown as dashed black lines. **i,** Estimation of the number of ScFc molecules per VLP based on the FCS molecular brightness analysis, for scFc-VLPs generated using scFc concentrations of 0.05 µM (s1), 0.1 µM (s2), 0.2 µM (s3), 0.5 µM (s4), and 1 µM (s5) (n=1).

To predict the scFc structure, we employed protein modeling approaches including AlphaFold2 (AF2, from DeepMind)^16^ and Rosetta^17^. The results showed that the Fc domain of the protein assumes conformations closely resembling the native Fc region **(Fig 1b)**. The presence of flexible linkers between the stably folded structural SNAPtag and scFc domains induces conformational dynamics. Protein conformational ensemble modelling showed that approximately 50-80% of the conformational states assumed by the SNAPtag are compatible with FcγR binding, with variability depending on the stringency of the applied threshold on allowed atomic contacts **(Extended Data Fig 1a)**. Additionally, the model allowed us to estimate the mean interdomain distance between the SNAPtag and Fc domain, falling in the range of 9.7 ± 0.9 nm (**Extended Data Fig 1b**).

The scFc was expressed in mammalian cells, ensuring the incorporation of glycan post-translational modifications on key residues that fine-tune the interactions between the Fc domain and Fc*γ*Rs.^18^ Non-reducing SDS-PAGE revealed the presence of monomeric and dimeric forms of the purified protein (**Extended Data Fig 2a**). To isolate the monomeric scFc, size-exclusion chromatography (SEC) was performed (**Extended Data Fig 2b**).

FcγR family members have different affinities for human IgGs.^1,19^ To characterize the binding properties of the scFc to key FcγRs, we immobilized the extracellular domains (ECDs) of FcγRIIa, FcγRIIb, and FcγRIIIa onto activated dextran surfaces of surface plasmon resonance (SPR) chips. The protein bound to all tested ECDs, despite the lack of Fab domains. The measured dissociation constants (K_D_) fell within the low micromolar range, and distinct binding kinetics for each FcγR class reflected differences in both association (k_on_) and dissociation (k_off_) rate constants **(Fig 1c)**. Importantly, these kinetic parameters were consistent with the reported parameters for the binding interactions between IgG and FcγRs,^19–21^ indicating that the design of the engineered scFc did not lead to impaired receptor binding.

### Multivalent assembly of scFc on a protein nanoparticle platform

We selected the well-characterized Hepatitis B core antigen (HBc) virus-like particle (VLP) as a protein display platform for precise, unidirectional, and multivalent presentation of scFc (scFc-VLPs).^22^ Through genetic fusion of the 12 kDa ΔN1SpyCatcher variant to the c/e1 loop within the major immunogenic region (MIR) of the HBc monomer (**Extended Data Fig 3a**), we introduced a SpyCatcher protein at the tip of the HBc spike. This SpyCatcher covalently binds the SpyTag scFc,^23^ displaying the protein on the surface of the VLP **(Fig 1d)**. The site-specific coupling enables the unidirectional display of the scFc.

The SpyCatcher-HBc monomers, expressed in *E. coli*, spontaneously self-assembled into VLPs which were then purified by an optimized disassembly-reassembly process (**Extended Data Fig 3b**-e). The process yielded particles with minimal contamination from single monomers (**Extended Data Fig 3f**) and uniform icosahedral shape with diameter of 30 nm, as validated by transmission electron microscopy (TEM) (**Extended Data Fig 3g**).

We generated a series of scFc-VLPs with different scFc valencies by varying the scFc concentrations (shown in parenthesis), while maintaining the VLP concentration constant at 4 *µ*M: s1 (0.05 *µ*M), s2 (0.1 *µ*M), s3 (0.2 *µ*M), s4 (0.5 *µ*M), and s5 (1 *µ*M) **(Fig 1e)**. The conjugation reaction reached saturation within 48 hours and yielded >90% display efficiency for all samples, except for s5 **(Fig 1f)**. The structural integrity of the particles was preserved after the scFc conjugation, with no aggregation observed by TEM (**Extended Data Fig 3g**). Furthermore, all scFc-VLPs exhibited uniform distribution of hydrodynamic diameters (D*_H_*), which increased with scFc concentration, as measured by dynamic light scattering (DLS) **(Fig 1g)**. We also observed an increase in the decay time as a function of scFc concentration (**Extended Data Fig 3h**), suggesting increased scFc valency on the VLPs.

To estimate the number of scFc molecules displayed on each scFc-VLP variant, we employed Fluorescence Correlation Spectroscopy (FCS), a biophysical method commonly used to characterize the diffusion properties and brightness of fluorescently labeled molecules or particles.^24,25^ We first fluorescently labeled scFc by a site-specific covalent reaction between the SNAPtag present in the scFc and ATTO 488-BG (**Extended Data Fig 4a**-c). The resulting scFc-488 exhibited physical stability in solution as confirmed by FCS (**Extended Data Fig 4d**,e). The scFc-488 was subsequently used to generate fluorescently labeled scFc-VLPs (**Extended Data Fig 4c)**. FCS measurements and brightness analysis revealed that the number of scFc molecules displayed on the VLP surface increased as a function of the scFc concentration **(Fig 1h,i)**. We estimated the number of scFc per VLP to range from 4 molecules for s1 to 59 molecules for s5 **(Fig 1i)** In addition, consistent with the DLS measurements **(Fig 1g)**, we observed an increase in the D*_H_* with increasing concentrations of scFc in scFc-VLPs (**Extended Data Fig 4f)**.

To elucidate the structural details of VLP and scFc-VLP, we conducted single-particle reconstruction using cryo-electron microscopy (cryo-EM) (**Fig 2, Extended Data** Fig 5). The 2D classification of the VLP demonstrated reasonable structural uniformity, with selected 2D averages exhibiting an inner mesh-like architecture and an outer amorphous layer formed by numerous protruding spikes (**Fig 2a**). The VLP reconstruction, with an overall resolution of 3.8 Å (gold-standard, FCS 0.143), revealed a capsid structure with icosahedron symmetry and a diameter of approximately 35 nm (**Fig 2b**). While the structural features of the inner shell domain were well-resolved, the outer surface layer displayed diffuse and poorly defined electron density, suggesting the presence of protruding SpyCatcher domains. The observed low local resolution is attributed to the inherent flexibility of the linkers used for the genetic fusion of the SpyCatcher domain to the HBc MIR. Structural alignment of the Hepatitis B virus (HBV) map (based on PDB 6BVN crystal structure) to the experimental VLP map confirmed a T=3 conformation. This conformation implies that the VLP is composed of 180 monomers forming dimers along distinct icosahedral axes, resulting in 90 spikes protruding from the VLP surface (**Fig 2c**). Next, we overlaid SpyCatcher-HBc structural conformers onto the HBV map, generating full capsid VLP predictive models **(Fig 2d)**. Computational analysis of the models disclosed an outer particle diameter (38.4 ± 1.6 nm), demonstrating variability due to diverse linkers conformations, and a mean minimal distance between SpyCatcher domains of 3.3 nm (**Fig 2e**). Using the HBV radial density profile alongside the radii of the VLP predictive models, we estimated the protrusion distance of the SpyCatcher domain from the VLP surface to be 4.5 ± 0.9 nm (**Extended Data** Fig 6). In addition, we conducted a reconstruction of the scFc-VLP, revealing an icosahedral structure with a diameter estimated to be 46 nm (**Fig 2f,g**). Interestingly, we observed an 11 nm internal shift compared to the VLP, potentially attributed to the presence of scFc molecules conjugated to the VLP surface. Similar to the VLP reconstruction, the density surrounding the inner shell domain of the scFc-VLP showed poor resolution due to the intrinsic flexibility of both scFc and SpyCatcher domains (Fig **2b,g**).

**Fig 2:**
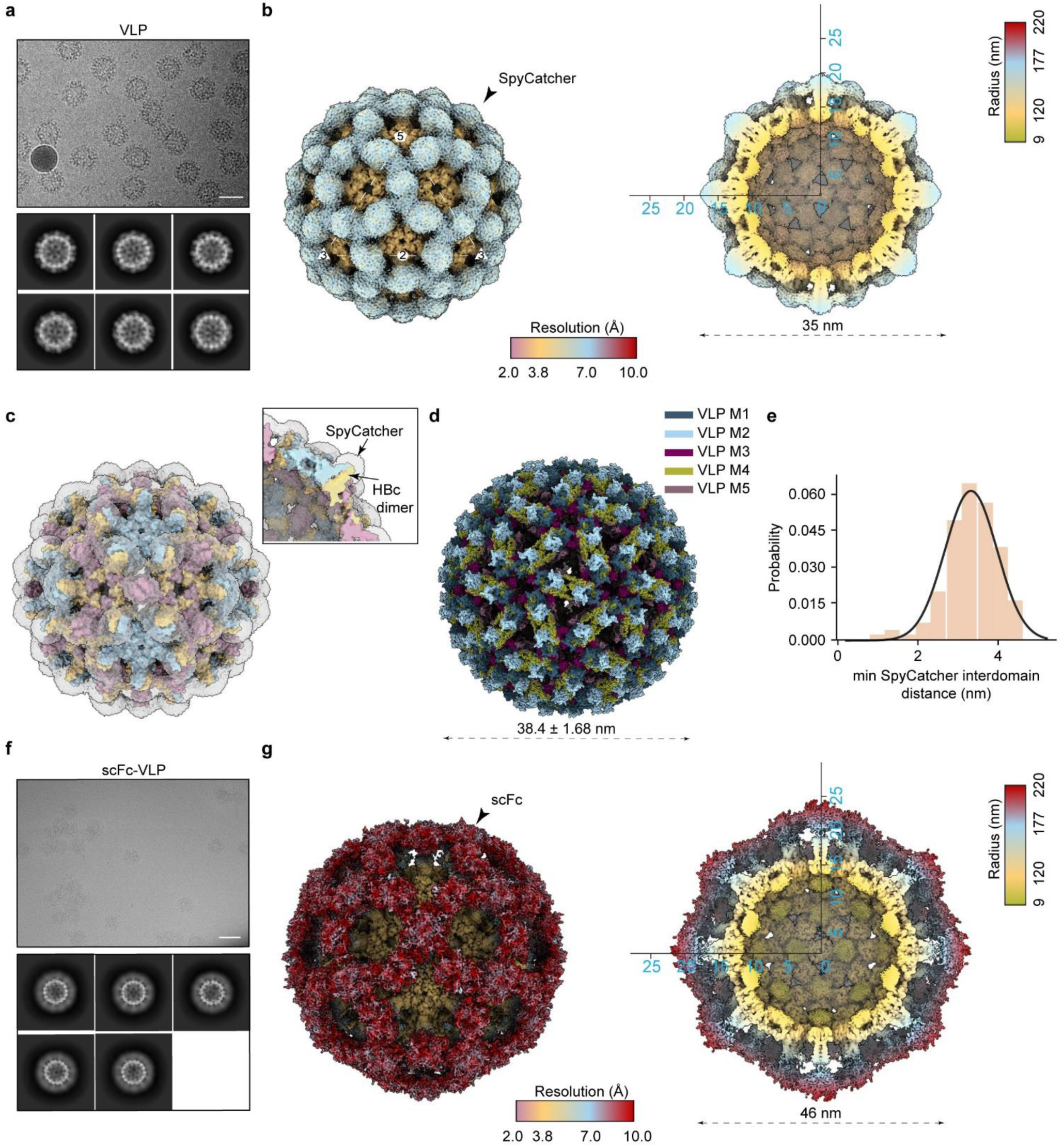
Structural characterization of VLP and scFc-VLP. **a,** Representative raw cryo-EM micrographs (top) and 2D-class averages (bottom) of VLP. Scale bar: 50 nm. **b,** Cryo-EM reconstruction of VLP (left panel), colored based on local resolution, as per the horizontal color bar (Å). Icosahedral axes (2-fold oval, 3-fold triangle, and 5-fold pentagon) are indicated. The low electron density at the tips of the VLP spikes suggests the presence of SpyCatcher domains. Bottom half of the VLP density map (right panel), radially colored. The calculated particle diameter is estimated to be 35 nm. **c,** Cryo-EM VLP map (depicted in transparent gray) overlaid with surface representation of HBV map. The HBV map is generated from HBV crystal structure (PDB 6BVN, 15 Å) assembled from an asymmetric unit composed of three identical chains (illustrated in red, blue, and yellow). The inset features a cross-section of the overlay, highlighting the alignment of the core structural elements (HBc dimer) between the two maps and the anticipated SpyCatcher domain, present in the VLP but not in the HBV map. **d**, The top five ranked VLP predictive models (VLP M1:5) are superimposed. Due to flexibility of the SpyCatcher linkers, used for the genetic fusion of the domains to the HBc MIR, the computed diameter of the particles range between 35.8 and 40.6 nm. **e**, Distribution of minimal distances between the center of mass of the SpyCatcher domains, computed based on the predictive structural models. The mean distance is estimated to be 3.3 nm. **f,** Representative raw cryo-EM micrographs (top) and corresponding 2D-class averages (bottom) of scFc-VLP. Scale bar: 50 nm**. g,** Cryo-EM reconstruction of scFc-VLP (left panel), colored based on local resolution, as per the horizontal color bar (Å). The low density surrounding the outer layer of the particle is presumed to correspond to the displayed scFcs on the surface of the VLP. Bottom half (right panel) of the scFc-VLP is radially colored. Estimated diameter of the scFc-VLP is around 46 nm.

Together, these results confirmed the formation of scFc-VLPs with a tailored number of scFc molecules per VLP. The cryo-EM reconstruction and FCS estimation revealed that VLPs can display from 4 to 59 scFc per particle, with the latter corresponding to scFc occupancy on almost two-thirds of the total VLP spikes.

### scFc valency on scFc-VLPs determines Fc**γ**R binding avidity

We employed SPR to study the binding kinetics of scFc-VLPs to low-affinity FcγRs (FcγRIIa, FcγRIIb, FcγRIIIa) and to C1q, an essential component of the complement cascade.^26–28^ VLPs did not bind to any of the immobilized FcγRs on the surface of SPR chips, whereas scFc-VLPs exhibited distinct binding profiles for each tested receptor **(Fig 3a-c**). A positive correlation emerged between scFc-VLP binding affinity and scFc valency on the VLPs, in line with previous studies showing the impact of Fc density on FcγR binding.^20^ Additionally, we also explored the effects of FcγR density on scFc-VLP binding. We observed that increasing the levels of immobilized receptors led to increased scFc-VLP binding. Importantly, the correlation between scFc valency and binding affinity was preserved (**Extended Data Fig 7a**-d), underscoring the robustness of these interactions.

**Fig. 3:**
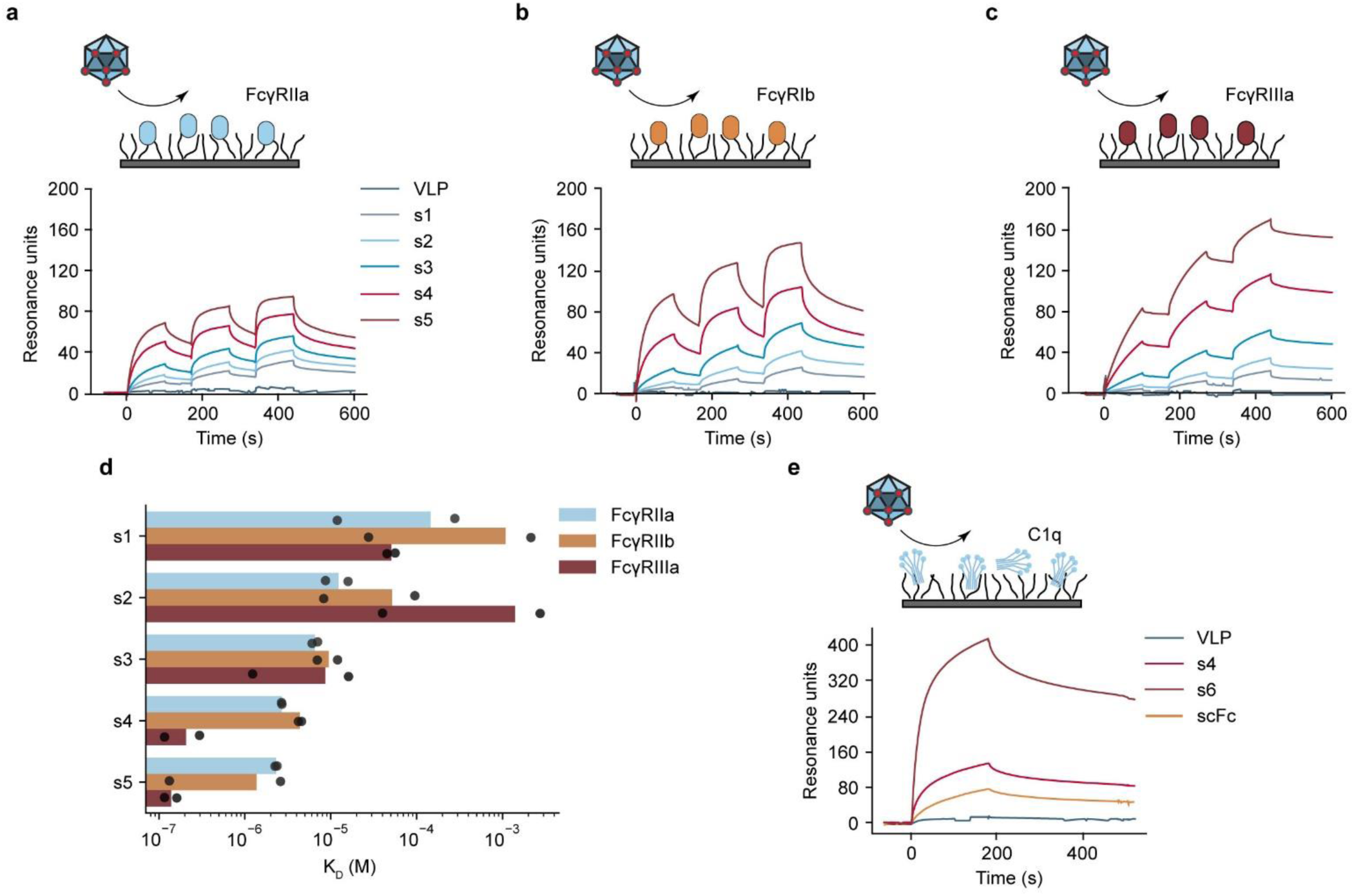
Valency of scFc in scFc-VLPs determines binding to FcγRs. **a-c,** Schematics of the SPR assay designs and sensorgrams. **a,** ECD-FcγRIIa, **b,** ECD-FcγRIIb, and **c,** ECD-FcγRIIIa were covalently attached to an activated dextran matrix and binding of VLP and scFc-VLPs (s1, s2, s3, s4, and s5) was assessed using single-cycle kinetic analysis. The analyte was injected in three increasing concentrations (0.15, 0.44, and 1.3 µM). One representative example of two independent experiments is shown. **d,** Comparison of K_D_ values of scFc-VLP binding to the different FcγRs tested, estimated by bivalent analyte model fitted to data shown in **a-c** (n=2). **e,** Schematic of SPR assay and SPR traces of binding of VLP, scFc, and scFc-VLPs (s4 and s6) to human C1q complex (n=1). For the construction of s6 we used VLP at 4 µM and scFc at 2 µM.

SPR analysis showed decreased K_D_ with increasing scFc valency on the VLPs for all receptors tested (**Fig 3d**), suggesting avidity effects. Further investigation using a dextran-free C1 chip, designed for studying large macromolecules and eliminating potential mass transport effects,^29^ demonstrated negligible binding for VLPs and scFc, while s3, s4, and s5 exhibited significant binding (**Extended Data Fig 7e**). These results confirmed the establishment of multivalent interactions between scFc-VLPs and FcγRIIa immobilized on the SPR surface.

Next, we studied the interaction between scFc-VLPs and C1q, which share overlapping FcγRs sites. While VLPs exhibited no binding to C1q, both scFc and s4 were able to bind C1q, with s4 showing a slightly higher binding affinity than scFc (**Fig 3e**). To validate the assay, we constructed a scFc-VLP variant with high scFc valency, named s6, given that C1q binding is known to depend on Fc density and organization. S6 demonstrated an approximately fourfold increase in binding compared with s4, which we attribute to the dense packing of scFc on the VLP surface, consistent with previous studies showing that high binding affinity and activation of C1q requires Fc clustering.^30^

In summary, our results highlight the specific and diverse binding capabilities of scFc-VLPs to FcγRs and C1q. The enhanced FcγR binding to s4 and s5, compared with scFc-VLPs with lower valencies, mirrors the increased receptor binding to multivalent IgG complexes compared with monomeric IgG.^3^ Furthermore, our findings underscore the potential of VLPs as a versatile platform for achieving precise and controlled multivalent binding of biomolecules.

### scFc valency on scFc-VLPs determines binding to immune cells

To analyze the interaction patterns of scFc-VLPs with immune cells, characterized by diverse FcγR expression profiles, we performed flow cytometry analysis of peripheral blood mononuclear cells (PBMCs) (**Fig 4a-d**, **Supplementary** Fig 1). In these experiments, VLPs and scFc-VLPs were labeled with GFP to enable their detection, and cells treated with GFP or PBS were used as controls. CD14+ monocytes, which express FcγRI, FcγRIIa, and to a limited extent FcγRIIb,^31^ exhibited significantly enhanced binding to all scFc-VLPs tested (s2, s4, and s5) compared with the controls (**Fig 4a**). FcγRIIIa-expressing natural killer (NK) cells^32^ showed specific labeling with s4 and s5 (**Fig 4b**), which was significantly increased compared with the controls and the s2. The specificity of the interactions was confirmed by the absence of scFc-VLP labeling of T cells, which lack FcγR expression (**Fig 4c**).^33^ scFc-VLPs and VLPs exhibited similar levels of binding to FcγRIIb-expressing B cells, which was therefore not specific (**Fig 4d**). Together, these results are consistent with the SPR data (**Fig 3a-c**), as the nanoparticles with highest scFc valency, s4 and s5, labeled monocytes and NK cells stronger than s2 or the controls. However, s5 did not result in higher cell labeling compared with s4 suggesting it did not exhibit increased binding to the FcγRs (**Fig 3d**). These findings show that the scFc-VLP binding to immune cells are cell type specific, indicating that not only receptor expression patterns but also the spatial organization of scFcs and FcγRs likely impact the binding efficiency of scFc-VLPs to immune cells.^34^

**Fig. 4:**
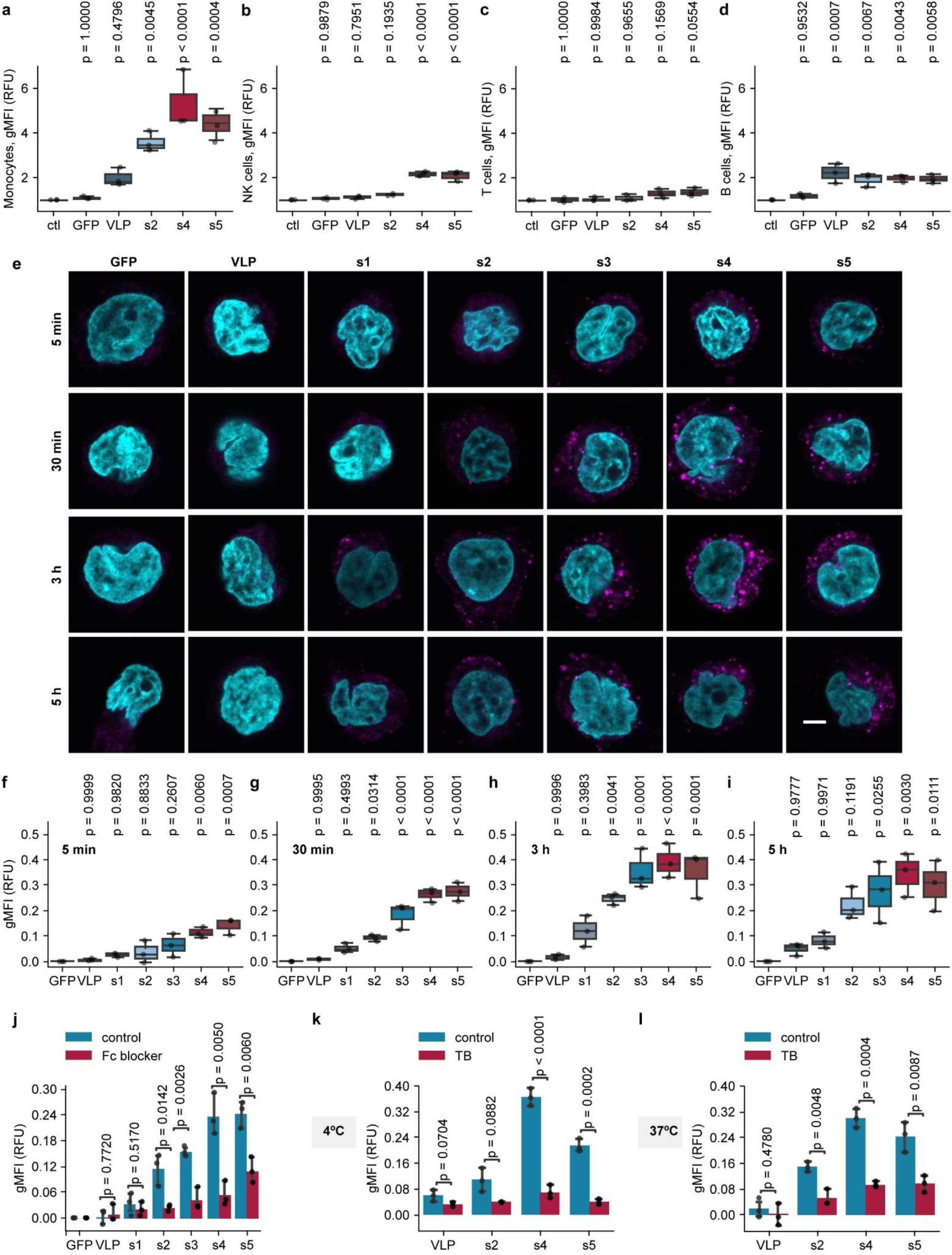
scFc valency controls scFc-VLP binding to immune cells. **a-d**, PBMCs were treated with GFP-labeled scFc-VLPs (s2, s4, s5), VLPs, or GFP for 30 minutes. Untreated cells served as a control (ctl). Quantification of the GFP signals by flow cytometry analysis in immune profiled PBMCs: **a**, monocytes; **b**, NK cells; **c**, T cells; **d**, B cells. gMFI, geometric mean fluorescence intensity. **e,** Confocal microscopy images of representative THP-1 cells treated with GFP-labeled VLP and scFc-VLPs (s1, s2, s3, s4, and s5), or GFP for the specified durations. Cell nuclei, stained with DAPI, are depicted in cyan, the GFP signal is shown in magenta. Each image is a representative of at least ten images from n=3 independent experiments. Scale bar: 5 µm. **f-i,** Time-resolved flow cytometry analysis of THP-1 cells treated with GFP-labeled VLP, scFc-VLPs (s1, s2, s3, s4, and s5), or GFP over specified durations: **f**, 5 min; **g**, 30 min; **h**, 3 hours; **i**, 5 hours. Untreated cells served as a control. **j,** THP-1 cells were preincubated with Fc blocker (red) or left untreated (blue) for 30 min, followed by addition of GFP-labeled scFc-VLPs (s2, s4, s5) or VLPs for 30 min. Untreated cells served as a control. **k** and **l,** THP-1 cells were treated with GFP-labeled VLPs and scFc-VLPs (s2, s4, s5) for 30 minutes, washed with PBS, and incubated with TB or left untreated (control). The GFP signal was measured shortly after the TB addition. Experiments were performed at 4°C (**k**) or 37°C (**l)**. TB, Trypan Blue. Data in all plots are presented as median ± std (n=3 independent experiments). Statistical analysis was performed using one-way ANOVA with Tukey’s multiple comparisons post-hoc test. P-values between the control and the indicated groups are shown.

To investigate the binding dynamics of scFc-VLPs to THP-1 cells, a human monocytic cell line that expresses high levels of FcγRII and FcγRIII,^35^ we used confocal microscopy (**Fig 4e**) and flow cytometry (**Fig 4f-i**). We detected a time-dependent increase in scFc-VLP binding to THP-1 cells, reaching saturation within 3 hours. THP-1 labeling increased with scFc valency on scFc-VLPs, in agreement with the results for monocyte staining in PBMCs (**Fig 4a**). The absence of nonspecific binding of VLPs and GFP to THP-1 cells indicated that the observed interactions were FcγR-mediated. To further confirm these results, we used a polyclonal FcγR-specific antibody to block Fc-mediated interactions in THP-1. In the presence of the Fc blocker, the levels of cell binding of all scFc-VLPs significantly decreased, with only minor nonspecific labeling observed for s5 (**Fig 4j**).

We employed Trypan Blue (TB) to quench extracellular fluorescence, thereby enabling differentiation between internalized and membrane-bound scFc-VLPs.^36^ In cells incubated with scFc-VLPs at 4°C, which constrains cell internalization, TB treatment led to a substantial decrease in the GFP fluorescence (**Fig 4k**). This effect persisted in THP-1 cells incubated at 37°C, indicating that a significant fraction of scFc-VLPs remained bound to the plasma membrane after 30 minutes (**Fig 4l**) and 3 hours (**Extended Data Fig 8a**), with s5 exhibiting slightly elevated levels of internalization. As a control, we performed the TB assay with FITC-IgG coated beads. In contrast to the scFc-VLPs, after 30 minutes of incubation, the particles were completely internalized (**Extended Data Fig 8b**,c).

In summary, our findings reveal that scFc-VLPs bind monocytes in a scFc valency-dependent manner yet exhibit negligible internalization. It has been previously shown that the structural properties of antibody bound epitopes can impact FcγR effector functions, with antigens exceeding 10 nm in length being able to abrogate the phagocytic process.^37,38^ The spikes protruding from the surface of scFc-VLPs, which have dimensions corresponding to the SpyCatcher domain protrusion (4.5 ± 0.9 nm) plus the predicted scFc length (9.7 ± 0.6 nm), impose structural constraints on scFcs displayed on VLPs, which could potentially interfere with FcγR activation. In addition, the fixed spatial arrangements of scFcs on the VLP surface can result in spatial organizations of scFc/ FcγR interactions that do not promote receptor activation.^39^

### scFc valency modulates FcγRII nanoscale organization and FcγRII activation in THP-1 cells

Using confocal microscopy, we investigated the spatial organization of FcγRII in THP-1 cells treated with IgG, *in vitro* assembled immune complexes (IC) or heat-aggregated IgG (HAIgG) and observed distinct receptor organizations depending on Fc valency (**Fig 5a**), in line with previous studies.^4^ IC and HAIgG, which are different forms of multivalent IgG, induced receptor aggregation, confirming the impact of avidity effects on receptor clustering.^40^ In contrast, monovalent IgG displayed an even distribution of FcγRII across the plasma membrane. Despite their multivalent scFc presentation, most scFc-VLPs (s1, s2, s3, and s4) did not induce FcγRII clustering. Occasionally, s5 induced the formation of FcγRII aggregates, which were smaller than the aggregates induced by HAIgG or IC treatment. Surprisingly, scFc alone induced significant receptor clustering, indicating that the unconjugated scFc acts as an FcγRII agonist.^40^

**Fig. 5:**
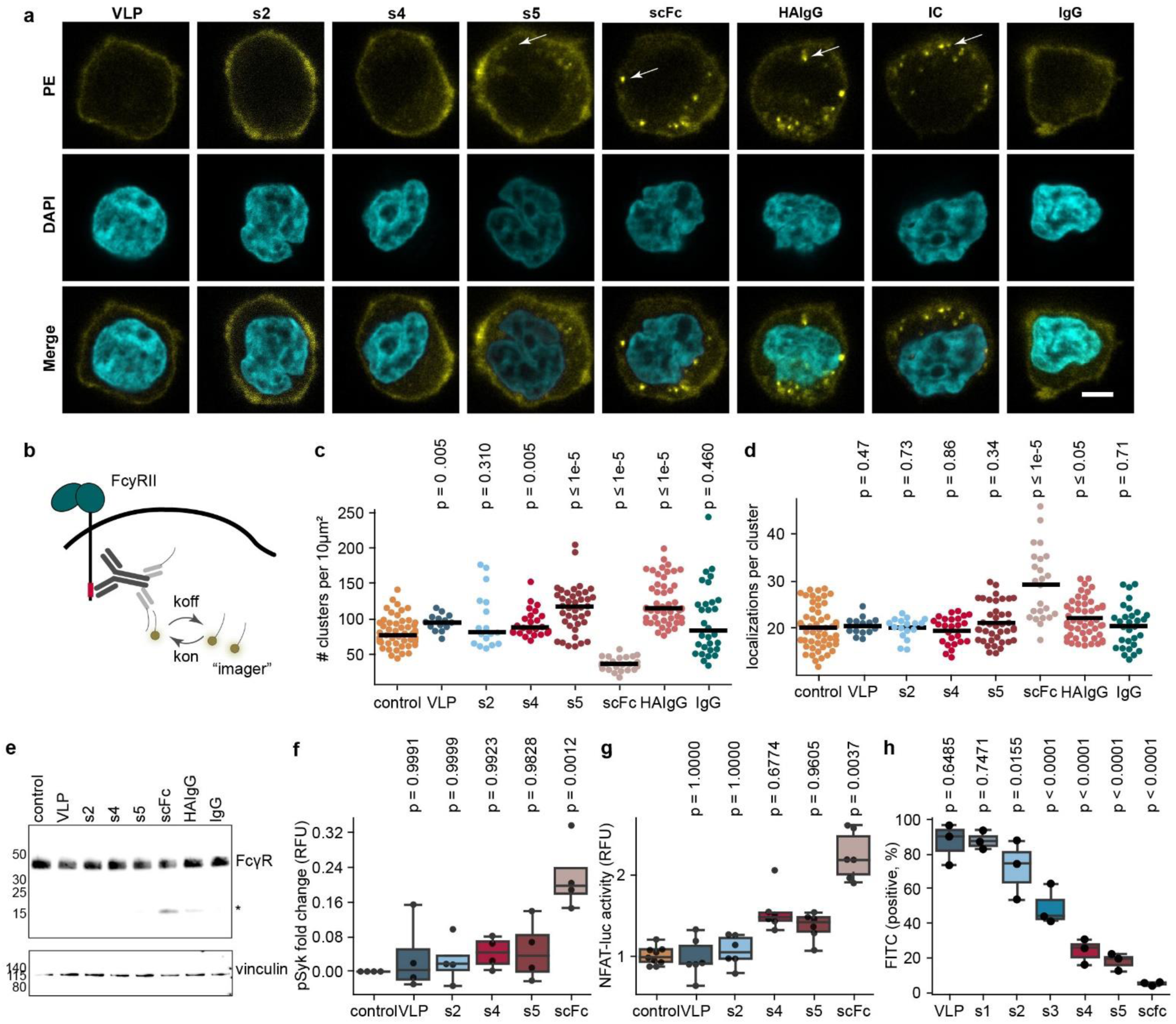
scFc but not scFc-VLPs induced oligomerization and activation of FcγRs. **a,** Confocal microscopy images of FcγRII-stained THP-1 cells treated with VLP, scFc-VLPs (s2, s4, s5), and scFc for 30 min at 37°C. HAIgG, IC, or IgG treatment served as controls. Arrows indicate large receptor aggregates. Each image is representative of at least 10 images from n=3 independent experiments. Cyan: DAPI stained cell nuclei; Yellow: FcγRII stained with PE-labeled anti-FcγRII antibody. Scale bar: 5 µm. **b,** Schematic of DNA-PAINT method used to image FcγRIIa in THP-1 cells. The DNA-PAINT signal is generated from transient interactions between short, fluorescently labeled DNA probes and complementary strands conjugated to secondary nanobodies bound to a FcγRII specific antibody (one or two nanobodies can bind to one FcγRII antibody). **c,** Number of detected clusters per 10 µm^2^ detected in THP-1cells left untreated (control) or treated with VLP, scFc-VLPs (s2, s4, s5), scFc, HAIgG, IC, or IgG for 30 min at 37°C. Horizontal lines indicate the median for each condition. P-values determined by a two-tailed Mann– Whitney test. **d,** Number of localizations per cluster detected in THP-1cells left untreated (control) or treated with VLP, scFc-VLPs (s2, s4, s5), scFc, HAIgG, IC, or IgG for 30 min at 37°C. Horizontal lines indicate the median for each condition. **e,** Immunoblotting analysis of FcγRII in THP-1 cells treated with VLP, scFc-VLPs (s2, s4, s5), scFc, HAIgG, and IgG for 30 min at 37°C. Untreated cells serve as controls. Cleaved FcγRII product denoted by *. The gel is representative of n=3 independent experiments. **f,** Phospho-flow cytometry of THP-1 cells for pSyk after a 3 min treatment with VLP, scFc-VLPs (s2, s4, s5), and scFc. Untreated cells served as a control (n=4, median ± std). **g,** NFAT activation in FcγRII Jurkat NFAT-luc reporter cells stimulated with VLP, scFc-VLPs (s2, s4, s5) and scFc. Luciferase activity measured after 8 h incubation. Untreated cells used as a control (n=3, median ± std). **h,** Percentage of FITC-labeled IgG-coated latex beads phagocytosed by THP-1 cells after 30-min treatment with VLP, scFc-VLPs (s1, s2, s3, s4, s5), and scFc (n=3, median ± std). Statistical analysis for **c** and **d** was performed by two-tailed Mann-Whitney test and statistical analysis for **f-h** was performed using one-way ANOVA with Tukey’s multiple comparisons post-hoc test. P-values calculated based on the comparison between the control and the indicated group.

To investigate FcγRIIa nanoscale organization at the cell membrane, we used DNA Point Accumulation in Nanoscale Topography (DNA-PAINT), a single-molecule localization microscopy method.^41,42^ DNA-PAINT enables precise spatial and quantitative assessments of proteins in close proximity to the plasma membrane (**Fig 5b, Extended Data Fig 9a**).^42,43^ We detected approximately 80 FcγRIIa clusters per 10 µm^2^ on the surface of THP-1 cells (**Fig 5c**), with an average radius of 40 nm (**Extended Data Fig 9b**). Quantitative analysis of localization clouds (**Extended Data** Fig. 9d-f) indicated that approximately 70% of the clusters were monomers and dimers, with the remaining forming small receptor islands (**Extended Data Fig 9g**) with nearest neighbor distances of approximately 249 nm (**Extended Data Fig 9h**). Following a 30-minute incubation with IgG, no alterations were observed in the spatial receptor organization and the cluster-defined parameters (**Extended Data Fig 9b**,c). Treatment with VLP and s4 induced a small, but significant, increase in the number of FcγRIIa clusters compared with the control. In contrast, HAIgG, s5, and scFc induced significant changes in receptor organization (**Fig 5c**), consistent with the confocal microscopy analysis (**Fig 5a**), which showed large-scale aggregation induced by these treatments. Intriguingly, scFc had the opposite effect on the number of FcγRIIa clusters compared with HAIgG and s5. Treatment with scFc resulted in a reduction of more than two-fold in the number of detected clusters (37 clusters per 10 µm²) (**Fig 5c**) but increased numbers of localizations per cluster (**Fig 5d**) and cluster density (**Extended Data Fig 9c**). In addition, scFc treatment resulted in a significant increase in the median distance between the edges of neighboring FcγRIIa oligomers, reaching 312.41 nm (**Extended Data Fig 9d**). Together, these findings suggest that scFc induced receptor internalization. In contrast, stimulation with HAIgG and s5 resulted in an increase in the number of detected clusters (110-120 clusters per 10 µm², **Fig 5c**) and increased cluster density (**Extended Data Fig 9c**), indicating retention of the receptor at the cell membrane. Notably, s4 did not affect the spatial organization of FcγRIIa, similarly to IgG and the control. These findings suggest that the structural context of scFc display modulates FcγRIIa nanoscale organization, with monovalent scFc inducing receptor oligomerization, unlike scFc presentation on VLPs.

To further explore the impact of scFc-VLPs on FcγRII, we employed immunoblotting. We failed to observe receptor degradation induced by scFc-VLPs (**Fig 5e**), in contrast to previous studies showing FcγRII degradation in response to treatment with Fc multimers.^44,45^ Instead, treatment with scFc and HAIgG resulted in site-specific receptor cleavage, evident by a cleaved product band in the scFc- and HAIgG-treated samples. This cleavage pattern resembles the loss of the Immunoreceptor tyrosine-based activation motif (ITAM) domain observed during FcγRII-mediated platelet activation.^46^ The findings align with the significant decrease in the number of FcγRII clusters in scFc-treated cells as shown by the DNA-PAINT analysis (**Fig 5c**).

In FcγRII-mediated signaling, receptor clustering is crucial for regulating receptor phosphorylation and activating key protein tyrosine kinases such as Syk, AKT, and ERK1/2.^47^ Previous studies have demonstrated that Fc valency and structural organization impact kinase activation. While monomeric and low valency Fc interactions induce minimal or no kinase phosphorylation, the multivalent Fc interactions drive receptor phosphorylation and downstream signaling.^4,45^ To evaluate the effects of scFc-VLPs on FcγR-mediated signaling in THP-1 cells, we first assessed Syk phosphorylation using flow cytometry (**Fig 5f**). scFc-VLPs did not induce kinase phosphorylation whereas scFc caused significantly increased Syk phosphorylation compared to controls. A cell-based assay using Jurkat T cells with an integrated NFAT luciferase reporter system and engineered to express FcγRII confirmed these results (**Fig 5g**). As FcγRs signal through the Syk-NFAT axis,^48^ luminescence serves as a readout for NFAT upregulation downstream of receptor activation. While scFc-VLPs showed no difference in luminescence signal compared with the untreated control, the scFc monomer caused a significant increase, about twice the level observed for the scFc-VLPs, indicating NFAT activation (**Fig 5g**). Together, these results show that scFc-VLPs have limited capacity to stimulate FcγR-mediated signaling in THP-1, in contrast to the effective activation induced by scFc.

Overall, these findings indicate that multivalent scFc display on VLP surface facilitates avid cell binding without inducing FcγRII membrane clustering as well as receptor-mediated signaling. This behavior is in stark contrast to the scFc monomer, which acts as FcγRII agonist.

### A high-valency scFc-VLP variant with low immunogenicity reduced IC-mediated phagocytosis *in vitro* and *in vivo*

Given the extensive labeling of FcγR-expressing monocytes by high-valency scFc-VLPs (s4 and s5) without inducing Syk activation, we investigated the capacity of these constructs to block FcγR functions. In an *in vitro* model of antibody-dependent cellular phagocytosis (ADCP) using FITC-labeled IgG-coated latex beads, THP-1 cells pre-treated with scFc-VLPs showed decreasing levels of phagocytosis with increasing the scFc valency on scFc-VLPs (**Fig 5h**). The VLPs did not interfere with the ADCP response. Interestingly, scFc exhibited an inhibitory effect on ADCP comparable to the high-valency nanoparticles.

To assess the ADCP-blocking efficacy of scFc-VLPs in a physiologically relevant context, we investigated antibody-mediated platelet phagocytosis – a hallmark of autoimmune disorders such as immune thrombocytopenia (ITP).^49^ Fluorescently labeled antibody-opsonized platelets were efficiently phagocytosed by THP-1 cells (**Fig 6a, Extended Data Fig 10a**), as previously reported.^50^ Treatment with scFc-VLPs led to a significant inhibition of platelet uptake, reflected in reduced THP-1 fluorescence, while VLPs had no impact on phagocytosis levels. Notably, the inhibitory effects of scFc-VLPs on platelet uptake were dependent on scFc valency, with high-decorated scFc-VLPs, s4 and s5, showing a statistically significant reduction in THP-1 fluorescence (**Fig 6a)**. Similar to the IgG-bead phagocytosis assay (**Fig 5h**), scFc showed potent FcγR inhibition. Together, these results demonstrated the ability of scFc-VLPs to interfere with the interaction of IC with FcγRs, suggesting that they have the potential to lead to therapeutic applications in autoimmune diseases characterized by autoantibody induction, such as ITP.

**Fig. 6:**
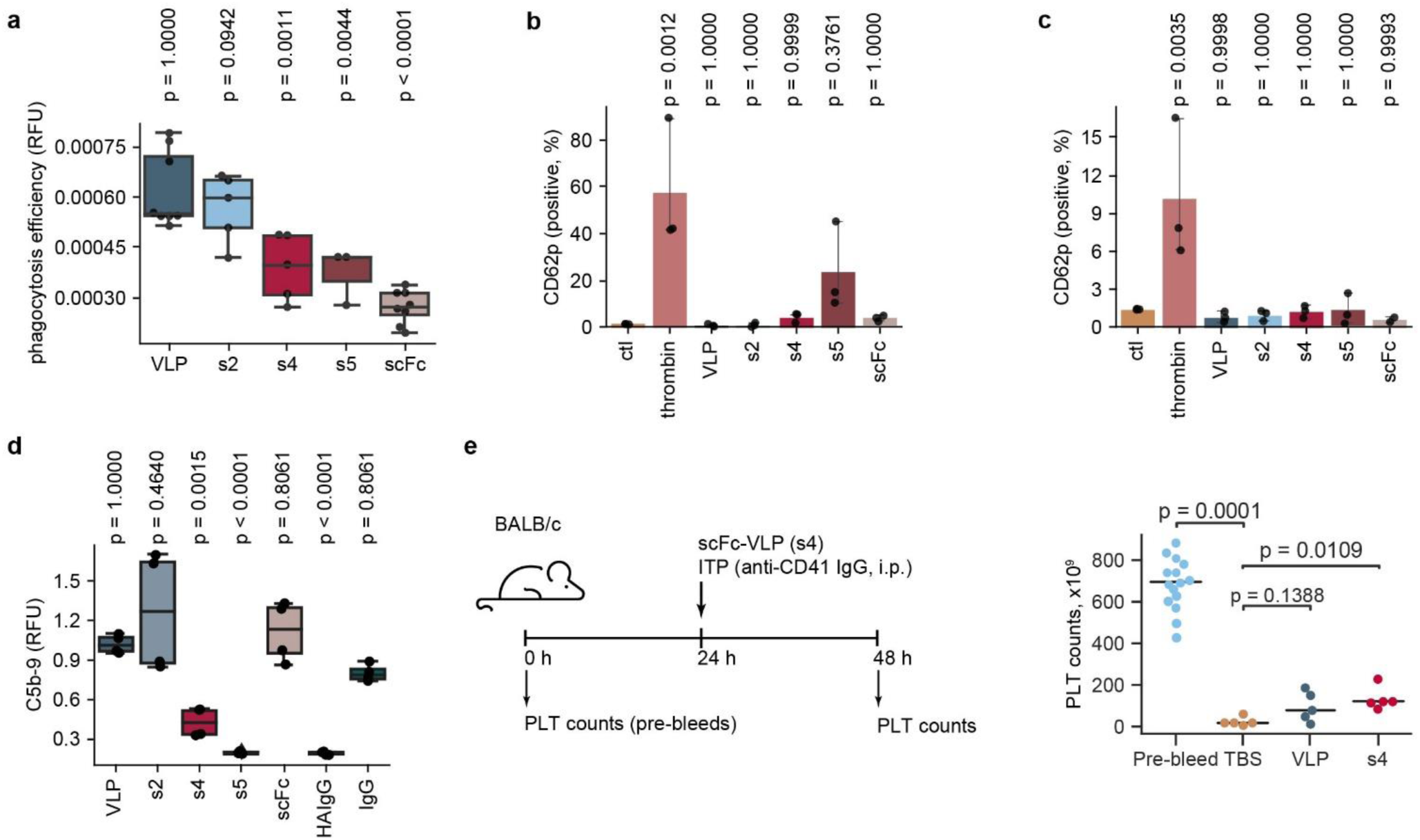
FcγRs blockade efficacy and immunogenicity of scFc-VLPs *in vitro* and *in vivo*. **a,** THP-1 cells pre-treated with VLP, scFc-VLPs (s2, s4, s5), scFc or PBS (control) were incubated with anti-HLA (W6/32) opsonized platelets. Phagocytosis efficiency was quantified by flow cytometry (n=5, mean ± std). **b** and **c,** Flow cytometry analysis of CD62p expression in human platelets treated with VLP, scFc-VLPs (s2, s4, s5), scFc, and thrombin (positive control) in the absence (**b**) or presence (**c**) of human plasma (n=3, mean ± std). **d,** Inhibition of the classical complement pathway by Normal Human Serum (NHS) using immobilized IgM as an activator, assessed via ELISA detecting soluble C5b-9. NHS was pre-incubated with VLP, scFc-VLPs, HAIgG, and IgG controls for 1 hour (n=3, mean ± std). Statistical analysis was performed using one-way ANOVA followed by Tukey post-hoc test for **a-d**. **e,** Platelet counts after treatment with s4, VLP or TBS (control) in ITP mouse model. Mean ± std are presented for biological replicates (n=5). Statistical differences between the anti-CD41 treated mice and the indicated group were determined using one-way ANOVA with Mann-Whitney multiple comparison test.

A critical safety consideration for *in vivo* applications of scFc-VLPs is the risk of platelet activation through FcγRIIa engagement. With FcγRIIa as the sole FcγR expressed in human platelets, the clustering and activation of platelet-FcγRIIa, indicated by elevated surface expression levels of CD62p (p-selectin), pose a potential threat of platelet aggregation, leading to the formation of microthrombi and life-threatening thrombosis.^51^ To assess the immunogenicity of scFc-VLPs, human platelets were exposed to the nanoparticles, and the surface levels of CD62p were measured. Among all tested conditions, only thrombin, used as a positive control, induced a significant increase in the CD62p+ platelet counts (**Fig 6b**). s5 caused a slight CD62p induction, which was not significant due to high donor-dependent variability. Additionally, in the presence of human plasma, platelet activation by scFc-VLPs was inhibited, likely due to FcγR blocking by endogenous human IgGs (**Fig 6c**). Furthermore, using mouse platelets lacking endogenous FcγRs, we observed increased CD62p expression only upon thrombin treatment (**Extended Data Fig 10b**,c).

Fc multimers exhibit the capacity to modulate complement system activation through stable interactions with C1q and other complement-associated proteins, leading to their functional disruption or sequestration from the cell surface.^2,5,44^ The valency-dependent binding of scFc-VLPs to surface-immobilized human C1q complex (**Fig** 3e) prompted us to assess their effects on the classical complement pathway (CP). To evaluate CP activation, we employed an enzyme immunoassay to specifically detect the formation of the terminal C5b-9 component. IgM precoated wells were incubated with normal human serum containing scFc-VLPs (**Fig 6d**). We observed a gradual increase in CP inhibition with an elevated Fc valency on the VLPs. Specifically, s2 showed no significant difference compared with the control, s4 exhibited potent inhibition, and s5 abolished complement activation with similar levels of C5b-9 compared with the HAIgG. In contrast, the VLP, scFc, and IgG control did not interfere with the CP.

Following the assessment of the functional and immunogenic properties of scFc-VLPs, s4 emerged as a promising candidate for subsequent *in vivo* evaluation. The construct demonstrated efficient FcγR blockade through high-avidity interactions (**Fig 4**), without inducing receptor activation (**Fig 5f**), and potent inhibition of FcγR-mediated phagocytosis (**Fig 6a**), and a favorable safety profile (**Fig 6b**).

To evaluate the *in vivo* efficacy of s4, we employed a murine model of passive ITP, a well-established model for therapeutic assessment. ITP was induced in BALB/c mice by intraperitoneal (i.p.) injection of rat anti-CD41 IgG1 antibody. After 24 h the platelet counts significantly decreased from 686 x 10^9^/L to 24 x 10^9^/L. In comparison to Tris-buffered saline (TBS), used as a negative control, s4 significantly increased the platelet counts in mice with ITP (**Fig 6e**). In contrast, VLP did not show a significant increase in the platelet counts compared with the control. We did not observe any systemic shock induced by the VLPs since the temperature of the animals was in the normal physiological after the i.p. injection (**Extended Data Fig 10d**). These results indicate that s4 has a positive impact on attenuating ITP and suggest that it holds promise as a design paradigm for the development of autoimmune therapeutics.

## Conclusion

The landscape of autoimmune disease therapeutics is on the verge of significant transformation, with Fc multimers emerging as promising candidates to overcome the limitations of the current standard, IVIG.^4,52^ The potential of Fc multimers lies in their ability to *en masse* block FcγRs, offering an effective alternative for treating IC-mediated autoimmune diseases. The efficacy of this approach relies on a delicate balance between achieving high receptor occupancy while preventing receptor activation. Advancing this therapeutic strategy requires an in-depth understanding of Fc-mediated FcγRs clustering dynamics and its correlation with receptor activation and induction of downstream signaling. To this end, it is necessary to understand the impact of various biophysical cues on FcγR activation, including Fc valency and spatial configuration.

In the field of Fc multimeric fusions, most presentations typically involve linear configurations with an undefined number of Fc units or non-linear forms with a constrained number of Fc domains – limited to a maximum of five Fc domains in branched (n = 1-5) or six Fc domains in hexameric structures. In this study, we employed a synthetic biology approach to display Fc domains on a protein-based platform with a tailored spatial organization. Through this approach, we achieved a considerable range in the number of Fc units, spanning from 4 to 59.

This engineered Fc display platform serves as a versatile tool for elucidating the impact of Fc valency as well as Fc structural context on FcγR clustering and activation. We observed a Fc valency-dependent increase in scFc-VLPs binding to diverse FcγR-expressing cells, displaying targeted specificity for monocytes and NK cells. In contrast to other Fc multimers,^4,44,45^ we observed low internalization and receptor clustering for the majority of scFc-VLPs, coupled with limited receptor activation. We hypothesize that the spatial context of the scFcs on the VLP platform determines the observed cellular responses. The geometry of scFc-modified VLP spikes, protruding more than 10 nm from the VLP surface, may constrain phagocytosis, as previously reported.^37,38^ Additionally, the fixed arrangement of the scFcs on the scFc-VLPs surface may lead to a spatial organization of scFc/FcγR interactions that are not conductive to FcγR activation.^39^

Together, our findings present a novel approach to suppress FcγR activation while blocking IC-mediated functions such as ADCP, thus offering insights into the potential of engineered Fc multimers for the development of innovative autoimmune disease therapeutics.

## Methods

### Construction, expression, and purification of scFc

For the construction of SNAPtag-scFc-SpyTag, hereafter referred to as scFc, two hIgG1 Fc domains, composed of CH2 and CH3 domains with preserved hinge region (GenBank accession number: AIC63048.1, spanning residues 177 to 407), were genetically fused together with a flexible (Gly_4_Ser)_3_ linker. A single cysteine to serine mutation (Cys_451_Ser) was introduced in the hinge region, disrupting one of the three disulfide bonds to increase flexibility.^53^ A SNAPtag (GenBank accession number: AQS79239.1) was fused directly to the N-terminus of the protein construct, and a SpyTag (AHIVMVDAYKPTK) was added to its C-terminus via a short GGS linker. The gene encoding the scFc was codon-optimized for mammalian expression, synthesized, and cloned into a modified pCDNA3.1(+) expression vector. A signal peptide (MEFGLSWLFLVAILKGVQC) optimized for expression of heavy antibody chains, was placed at the beginning of the scFc sequence to facilitate protein secretion.^54^ The plasmid was transfected into HEK 293F cells (Thermo Fisher Scientific). After 88 h of incubation, the cell suspension was harvested by centrifugation at 200 × g for 5 min. The supernatant was collected, cleared by centrifugation at 6000 × g for 10 min, and filtered through a Whatman Polycap 36TC filter. The sample was buffer-exchanged into Tris-Tween-20 buffer and concentrated using a 10 kDa MWCO membrane, followed by five rounds of buffer exchange into 20 mM H_2_PO_4_, pH 7.1. Next, the sample was loaded onto a 5 ml HiTrap Protein G HP column (Cytiva) at a flow rate of 5 ml/min. The elution was performed using 0.1 M Glycine at pH 2.7, with the collected fractions immediately neutralized using 1 M Tris, pH 9. Fractions corresponding to the main elution peak were pooled and concentrated using a 30 kDa MWCO Vivaspin20 filter device (Cytiva) with a swing-out rotor at 3900 × g for approximately 6 h. The protein sample was loaded onto a HiPrep 16/60 Sephacryl S-200 HR column (Cytiva) pre-equilibrated with 0.2 µm filtered and degassed PBS. A distinct peak in the chromatogram with a molecular mass of approximately 80 kDa was collected and concentrated using a 30 kDa MWCO Vivaspin20 filter device (Cytiva). The protein was stored at -20°C.

### Construction, expression, and purification of SpyCatcher-VLPs

SpyCatcher-HBc was constructed and cloned into a pET24a vector using standard PCR methods and Gibson isothermal assembly.^55^ ΔN1SpyCatcher (GenBank accession number: AFD50637.1, spanning residues 50– 129) was integrated into the surface-exposed major immunodominant loop of truncated HBc (GenBank accession number: BAG49610.1, spanning residues 1–149) between amino acids 78 and 81. Flexible linkers were placed at the N- and C-terminus of SpyCatcher to promote flexibility and hexahistidine tag (His6x) was introduced at the C-terminus. The insert was codon optimized for expression in *E. coli* and verified by Sanger sequencing. BL21(DE3) cells (Thermo Fisher Scientific) were heat-shock transformed with the SpyCatcher-HBc expression plasmid, plated on LB-agar (50 µg/mL chloramphenicol), and incubated for 16 h at 37°C. A single colony was selected and inoculated in 10 ml LB medium (50 µg/mL chloramphenicol) overnight, at 37°C, with continuous agitation at 220 rpm. The cell culture was diluted 1:100 into LB medium (50 µg/mL chloramphenicol) and incubated at 37°C, 220 rpm until reaching an optical density at 600 nm (A_600_) of 0.6. Protein expression was induced by adding 0.8 mM isopropyl β-D-1-thiogalactopyranoside (IPTG) and incubating for 16 h at 22°C, 180 rpm. Cells were harvested by centrifugation at 5000 **×** g for 15 min at 4°C and the cell pellet was resuspended in a lysis buffer composed of TN300 buffer (5 mM Tris-HCl, 300 mM NaCl, pH 8.5) supplemented with 0.1 mg/mL lysozyme, 1 mM phenylmethanesulfonyl fluoride (PMSF), 0.1% Triton X100, 0.1% Triton X114, and 0.1% Tween-20. The lysate was incubated for 30 min at RT on an orbital shaker and sonicated using a probe sonicator on ice four times for 60 s at 50% duty cycles. The supernatant was removed by centrifugation at 35000 × g for 45 min at 4°C. The insoluble fraction containing SpyCatcher-VLP was resuspended in a dissociation buffer (4 M urea, 200 mM NaCl, 50 mM sodium carbonate, 10 mM 2-mercaptoethanol, pH 9.5) and incubated overnight at 4°C on an orbital shaker. Next, the pellet was discarded by centrifugation at 18000 × g for 45 min at 4°C and the supernatant was filtered through 0.45 and 0.22 µm syringe filters. The SpyCatcher-HBc monomers were purified by Ni^2+^- chelate affinity chromatography. The sample was processed through a HisTrap FF column (Cytiva) using fast protein liquid chromatography (FPLC) with an ÄKTA pure FPLC system (Cytiva). The flow rate was set to 1 ml/min, and the column was washed with 30 column volumes of dissociation buffer and 20 column volumes of dissociation buffer supplemented with Triton X114. The column was kept at 4°C. The SpyCatcher-HBc monomers were eluted with a dissociation buffer containing 0.5 M imidazole. Elution fractions containing the protein were collected, pooled, and subjected to stepwise dialysis for further processing. Specifically, protein fractions were dialyzed against dialysis buffer 1 (0.5 M urea, 100 mM Tris, 150 mM NaCl, 2 mM DTT, 1 mM EDTA, 10 mM CaCl2, pH 8.0) using Slide-A-Lyzer™ MINI Dialysis Devices, 10 K MWCO at 4°C for 4 h, allowing the SpyCatcher-HBc protein to start self-assembling. Then, the solution was dialyzed against dialysis buffer 2 (100 mM Tris, 150 mM NaCl, 1 mM EDTA, 10 mM CaCl2, pH 8.0) to completely remove urea and allow DTT to completely reassemble the VLPs. For some experiments, the reassembled particles were filtered through 0.2 µm PES membrane (Agilent), and the sample was purified by SEC-HPLC on a Bio SEC-5, 2000 Å, 21.2 × 300 mm, 5 µm (Agilent). Next, the particles were centrifuged for 2 min at 2200 rpm to remove aggregates and the supernatant was collected and concentrated using a 100 kDa MWCO Vivaspin20 filter device (Cytiva). Endotoxin concentrations were determined with Pierce LAL Chromogenic Endotoxin Quantitation Kit (Thermo Fisher Scientific) following the manufacturer’s instructions and were below 1 endotoxin unit/mL for SpyCatcher-VLP preparations. Quantification of the VLPs was carried out using both Nanodrop and a BCA protein assay kit (Pierce) using a bovine serum albumin (BSA) standard curve.

### scFc and SpyCatcher-VLP monomer structural predictions

To access the impact of introducing SNAPtag, henceforth referred to as the 21–224 region, through a flexible linker on the scFc binding to the human FcγRIIa, the structure of the complex was predicted with AlphaFold2 (AF2). Residues 21–694 of the scFc construct in complex with FcγRIIa were submitted to local ColabFold v1.5.2^56^ with template functionality and AMBER relaxation enabled. The sequence of FcγRIIa was obtained from PDB entry 3RY6 chain C.^57^ AF2 failed to predict stereochemically meaningful conformations for the linker segments. To resolve energetic issues, the selected predicted model was energy refined using the FastRelax protocol^58^ from the Rosetta molecular modeling suite,^59^ starting from a model where a torsion value of the linker was adjusted to prevent the folded domains from clashing during the structural optimization trajectory. The model was restrained to input coordinates with the flag `-constrain_relax_to_native_coords` and the best model out of 100 simulations was selected. The energy-refined model was used as an input for a simulation of energetically relevant linker conformations. Following Potrzebowski et al.^60^ a Monte Carlo simulation of the linker torsion angles of the residues 191– 224 was run guided by the Rosetta energy function, using 1000 torsion moves per simulation. In total 10000 simulations were run and the 1000 models with the lowest energy were selected for analysis. To estimate the decrease in affinity to the Fc receptor wrought by the inclusion of the 21–224 region in the scFc construct, the fraction of models in the generated ensemble where the Fc receptor was obstructed was calculated. Whether the 21–224 region invaded the volume that the receptor occupies in complex with the scFc construct was measured by reintroducing the structure of the receptor in each computational model, optimizing of the positions of the side chains of the amino acids in the interface between the scFc construct and FcγRIIa, and then checking for clashes after. A clashing conformer was defined based on the difference between the `interchain_vdw` score term in Rosetta between the model in question and the Fc receptor in complex with the scFc construct with the 21–224 region deleted. The fraction of conformer clashing presented in the main text is the result of the two thresholds: a stringent threshold of 0 Rosetta energy units (REU) and a lenient threshold of 5 REU. The distribution of the distance between the scFc and SNAP-tag domains was estimated from the centers of mass (COM) and radii of gyration (RG). The interdomain distance was defined as the sum of the RGs of both domains and the distance between the COMs. The SpyCatcher-HBc dimer was similarly predicted using local ColabFold followed by FastRelax. To model alternative conformations of the SpyCatcher domain, a chain break was introduced between residue 184 and 185, cuting the glycine linker corresponding to residues 177–191, after which conformational variants of the other glycine linker at 79–93 were generated with Monte Carlo. The Monte Carlo simulation was run as described above, but with modeling with symmetry modelling enabled^61^ and using 25000 models. The cut glycine linker at 177–191 of the 5000 lowest-energy models were reconnected using Kinematic Loop Closure in Rosetta^62^ based on a monomeric subunit. The final ensemble was generated from the models with a ChainBreak score of < 0.02. An atomic model of HBV capsid was initially generated by VIPERdb^63^ applying icosahedral symmetry to an asymmetric subunit, consisting of three identical chains (A, B, C), each represented as 60 copies. The coordinates of the chains were obtained from PDB ID 6BVN.^57^ The Situs package^64^ was used to process the HBV crystal structure, generate density map and fit it to the experimental VLP map. The SpyCatcher-HBc monomers were superimposed upon the 180 subunits of the atomic structure of the T=3 HBV capsid (PDB ID 6BVN) to generate VLP capsid models. The minimal distance between the SpyCatcher domains displayed on the VLP capsid surface was estimated based on the COM of the SpyCatcher domains.

### scFc conjugation to VLP

SpyTag-scFc at concentration of 0.05, 0.1, 0.2, 0.5 or 1 µM was conjugated at 4°C with 4 µM SpyCatcher-VLP in TBS, pH 8.0. The reaction was incubated for different time periods ranging from 5 min to 72 h. The samples were mixed with reducing sample buffer (NuPAGE™ LDS Sample Buffer (4X), 5% beta-mercaptoethanol), followed by heating for 5 min at 95°C. Samples were analyzed by SDS-PAGE using 4-12% NuPAGE Bis-Tris gels (Thermo Fisher Scientific). The gels were stained with a silver staining kit (Pierce), according to the manufacturer’s instructions. The conjugation efficiency of SpyTag-scFc to SpyCatcher-HBc was calculated as 100 * (band intensity of conjugated scFc-VLP/(band intensity of conjugated scFc-VLP + band intensity of unconjugated scFc)).

### Dynamic light scattering (DLS)

VLP and scFc-VLPs were mounted using 1.0 × 1.0 mm disposable cuvette capillaries with a thickness of 200 *µ*m (Malvern, ZSU0003). The capillaries were placed within a low-volume disposable sizing cell Kit holder (Malvern, ZSU1002). Light scattering readings were measured using a ZetaSizer Ultra (Ultra ZS; Malvern). The data were obtained from three different measurements performed at 25°C.

### Transmission electron microscopy (TEM)

VLP and scFc-VLPs (4*µ*M) were applied onto a glow-discharged carbon-supported copper TEM grid (TEM-CF200CU50, Thermo Fisher Scientific) and incubated for 60 s before gently removing the solution. Subsequently, the grids were stained for 10 s with 5 *µ*l of 2 % w/v uranyl formate, which was carefully removed. This staining procedure was repeated seven times, followed by air-drying of the TEM grids for 30 min before imaging. The imaging process was conducted using a Talos 120C G2 (120 kV, Ceta-D detector) at × 92,000 x magnification for detailed near-field views. The acquired raw images were subjected to processing using ImageJ software (v1.53) for further analysis.

### Cryo-EM sample preparation and image processing

VLP (10 µM) and scFc-VLP (4 µM) were used and 3 µl sample aliquots were applied to CryoMatrix holey grids with amorphous alloy film (R 2/1 geometry; Zhenjiang Lehua Technology Co., Ltd), glow-discharged with 25 mA for 2 min using an EMS 100X (Electron Microscopy Sciences) glow-discharged unit prior to sample application. The grids were vitrified in a Vitrobot Mk IV (Thermo Fisher Scientific) at 4°C and 100 % humidity with 10 s blotting. Cryo-EM data collection was performed with EPU automated data acquisition software (Thermo Fisher Scientific) using a Krios G3i transmission-electron microscope (Thermo Fisher Scientific) operated at 300 kV acceleration voltage. During the exposure 60 (VLP) or 30 (scFc-VLP) movie frames were collected with a flux of 0.72 (VLP) and 1.01 (scFc-VLP) e-/ Å^2^ per frame. Raw images were motion corrected on the fly using Warp.^65^ All further processing steps were carried out using CryoSPARC v4.2.1.^66^ Initially aligned, non-dose-weighted micrographs were used to estimate the constant transfer function (CTF). Particles were selected using a template picker and then cleaned via two and three-dimensional classification. A reference model was generated using Ab-Initio Reconstruction with icosahedral symmetry restraints. Whole VLPs were reconstructed via homogeneous nonuniform refinement, with I1 symmetry imposed. All reported resolution values are a result of independent half maps analysis with gold standard FSC criterion (FSC = 0.143). All figures containing cryo-EM maps were prepared using ChimeraX. Cryo-EM collection parameters are presented in Supplementary Table S1.

### Fluorescence correlation spectroscopy (FCS)

The scFc protein was labeled with SNAP-Surface 488 (BG-488, Biolegend). In brief, BG-488 was dissolved in anhydrous DMSO and added to the scFc solution in a 1:5 molar excess, in the presence of reducing agent (1 mM DTT). The mixture was then incubated for 30 minutes at 37°C. To remove the unreacted dye, Zeba spin desalting columns (7K, Thermo Fisher Scientific) were utilized.

FCS measurements were performed on an FCS-equipped Zeiss LSM 980 confocal microscope at 25°C. 488 nm excitation of ATTO 488 yielded a diffusion time of 39 ms corresponding to a radius of the detection volume of ω=0.25 mm and a volume of 0.45 fL. Measurements were performed on #1.5 glass-bottom dishes (MatTek Corp.). FCS analysis was performed using the Zeiss Zen software and Microcal Origin (OriginLab). The autocorrelation curves were fitted with a one- or two-component model:

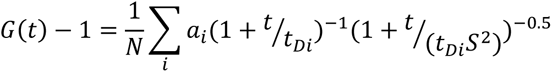

where N is the mean number of fluorescent molecules within the detection volume, τ_D_ is the diffusion time, S=z/w where z is the half-height and w is the radius of the detection volume. The confocal volume is given by:

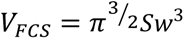

The relationship between the *τ*_D_ and the ω is given by:

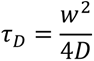

Stokes-Einstein equation was subsequently used to calculate hydrodynamic radii (*D_H_*) via the obtained diffusion coefficients (*D*). The number of proteins conjugated to the VLP was calculated by normalizing the counts per particle (CPP) of the scFc-VLP by the brightness of the free scFc.

### Surface plasmon resonance (SPR)

SPR experiments were conducted using a Biacore T200 instrument (Cytiva) equipped with Biacore T200 system control software v. 2.01. Recombinant human Fc*γ*IIa (Biolegend), Fc*γ*RIIb (Biolegend), Fc*γ*RIIIa (Biolegend), or human complement C1q (Sigma-Aldrich) were immobilized on a CM5 or C1 sensor chip (Cytiva) via amine coupling (Cytiva), following the manufacturer’s instructions. The scFc, scFc-VLPs, and VLP samples were injected at three increasing concentrations (0.15, 0.44, and 1.3 *µ*M) at a flow rate of 20 *µ*L/min in a single-cycle kinetics mode, or at one concentration (1.3 *µ*M) in a standard kinetic mode. HBS-P+ buffer (Cytiva) was used as the running buffer. Data were globally fitted to the bivalent analyte binding model and all kinetic parameters were determined by the BIAevaluation 3.0 software. The bivalent analyte model calculates separate kinetic constants for the affinity determining first step (*kd1*) and the avidity determining second step (*kd2*) of the binding process. We calculated the *K_D_* values based on the ratio between *kd_1_* and *ka_1_*.

### Cell lines and culture

THP-1 cells (German Collection of Microorganisms and Cell Cultures DMSZ) were maintained in Gibco™ RPMI 1640 Medium, GlutaMAX™ (Thermo Fisher Scientific) supplemented with 10% heat-inactivated fetal bovine serum (HI-FBS) (Sigma Aldrich) at 37°C, 5% CO_2_. Cells were split at a density of 1 x 10^6^ cells/ml.

### Preparation of heat-aggregated IgG (HAIgG)

IgG from human serum (Sigma-Aldrich) was incubated at 62°C for 30 min to induce aggregation. Subsequently, the sample was centrifuged at 13,000 x *g* for 10 min at 4°C to remove large-scale aggregates. The HAIgG was stored at 4°C and used within 24 h.

### scFc-VLP binding to PBMCs

PBMCs (BPS Bioscience) were resuspended in 10 ml AIM V™ medium (Thermo Fisher Scientific) with DNAse I (Sigma-Aldrich) and incubated for 30 min at 37°C, 5% CO_2_. The cells were centrifuged at 300 x g for 10 min and resuspended in assay medium [Gibco™ RPMI 1640 Medium, GlutaMAX™ (Sigma-Aldrich), 10% ultra-low IgG FBS (Thermo Fisher Scientific)]. Next, 0.2 x 10^6^ cells were left untreated or treated with scFc-VLPs labeled with 0.1 µM SpyTag-GFP, herein referred to as GFP, for 30 min at 37°C, 5% CO_2_. The cells were washed twice with a staining buffer [PBS, 0.5% BSA (Sigma-Aldrich), and 0.01% NaN3 (Sigma-Aldrich)] and labeled for 1 h on ice with an antibody lineage cocktail containing CD3-BV510 (UCHT1, Biolegend), CD14-BV421 (M5E2, Biolegend), CD19-APC (Thermo Scientific), CD56-PE-Cy7 (eBioscience™), and CD32-PE (Biolegend), along with Zombie NIR™ (Biolegend). The PBMCs were washed twice and analyzed with a BD FACSCanto II flow cytometer (BD Biosciences AG). Data analysis was performed using FlowJo 10.7.1 software. The degree of cell labelling was determined by normalizing the geometric mean of fluorescence intensity of each sample to the untreated control.

### scFc-VLP binding to THP-1 cells

THP-1 cells (0.2 x 10^6^) were suspended in assay medium and treated with 0.5 µM GFP-labeled VLP or scFc-VLPs, or 0.1 µM GFP alone. The samples were incubated for 30 min at 37°C, 5% CO_2_. For the Fc blocking assay, the cells were pre-treated with Fc receptor binding inhibitor polyclonal antibody (eBioscience™, Thermo Fisher Scientific) for 30 min. Next, the cells were washed with PBS to remove unbound probes, fixed in 4% paraformaldehyde in PBS (PFA/PBS) for 10 minutes, washed 2x with PBS, stained with DAPI, washed 2x with PBS. THP-1 labeling was quantified by flow cytometry using a BD FACSCanto II flow cytometer (BD Biosciences), and data analysis was conducted with FlowJo 10.7.1 software. For confocal microscopy, 50 µl cell suspension was seeded onto 0.1% gelatin-coated 18-well glass-bottom chambers (Ibidi) and incubated for at least 30 minutes at 4°C. Image acquisition was performed on a Zeiss LSM900-Airy2 scan confocal microscope with a 63X/1.4 oil objective. Images were processed using ImageJ and a custom-built Python script.

### scFc-VLP internalization assay

THP-1 cells (0.2 x 10^6^) were resuspended in assay medium and treated with 0.5 µM GFP-labeled VLP or scFc-VLPs, or 0.1 µM GFP alone, and incubated for 30 min or 3 h at 37°C, 5% CO_2_, or for 30 min at 4°C. Subsequently, cells were washed with PBS and placed on ice. Prior to the flow cytometry analysis of THP-1 labeling, each sample was divided, and 4 mg/ml Trypan Blue (Thermo Fisher Scientific) was added to one of the fractions to quench extracellular fluorescence. The flow cytometry quantification and analysis were performed as described before.

### THP-1 bead phagocytosis assay

THP-1 cells (0.2 x 10^6^) were resuspended in assay medium and treated with 0.5 µM VLP, scFc-VLPs, or 1 µM scFc for 30 min at 37°C, 5% CO_2_. Untreated cells were used as a control. Next, latex beads coated with FITC-IgG were added in 1:500 dilution and the samples were incubated for 30 min, 37°C, 5% CO_2_. Afterwards the cells were washed with PBS to remove unbound beads, fixed with 4% PFA/PBS for 10 min at 25°C and resuspended in assay buffer (Cayman Chemicals). Flow cytometry was performed using BD FACSCanto II flow cytometer (BD Biosciences AG) equipped with BD DIVA software and the analysis was performed with 10.7.1 software.

### THP-1 platelet phagocytosis assay

Whole blood was obtained from human volunteers via venipuncture with ethical approval from Etikprövningsmyndighetens (Reference: Dnr 2022-05723-01). The work is in accordance with the Code of Ethics of the World Medical Association (Declaration of Helsinki), written consent was acquired from all participants. The obtained whole blood was centrifuged at 180 x g for 10 min at 25°C, and the top layer, containing platelet-rich plasma (PRP), was carefully collected. PRP was washed in PBS and subsequently stained with pHrodo (Thermo Fisher Scientific, 40 µM) for 30 min at 25°C in the dark. Following staining, the platelets were washed twice in PBS and opsonized with anti-human HLA class I (R&D Systems, 2.5 µg/mL), IgG control (R&D Systems, 2.5 µg/mL), or PBS for 30 min at 25°C. THP-1 cells (0.1 x 10^6^ cells per sample) were washed, resuspended in RPMI (HyClone Laboratories) and stained with CellMask Deep Red (Thermo Fischer Scientific, 5 µg/mL) for 30 min at 25°C in the dark. Subsequently, the cells were washed and resuspended at 10^6^ cells/mL in RPMI supplemented with 0.5 µM VLP, 0.5 µM scFc-VLPs, 1 µM scFc, or with TBS for 30 min at 25°C. Stained platelets (10 x 10^6^) were mixed with 100 µL treated THP-1 and incubated at 37°C, 5% CO_2_ for 1 h. Finally, the cells were washed, resuspended in RPMI and subjected to flow cytometry analysis (Accuri C6, BD Biosciences). All washing steps were performed at 500 x g for 5 min at 25 °C. The phagocytosis efficiency was calculated by normalizing the signal from each condition by the signal from both positive control (THP-1 cells premixed with anti-human HLA class I opsonized platelets) and negative control (THP-1 cells premixed with mouse IgG opsonized platelets).

### Platelet activation assay

PRP was obtained by centrifuging human or mouse whole blood. Human whole blood was obtained as previously described. Mouse whole blood was centrifuged at 350 x g for 5 minutes, and the supernatant enriched with PRP was carefully collected. For the isolation of washed platelets, PRP was centrifuged at 500 x g for 5 min. The supernatant was discarded, and platelets were resuspended in an equal volume of PBS. Subsequently, PRP was incubated with GPRP (Sigma-Aldrich, 5mg/mL) for 1 min. Both the treated PRP and the washed platelets were treated with 0.5 µM VLP, 0.5 µM scFc-VLPs, 1 µM scFc, bovine thrombin (Sigma-Aldrich, 1 U/mL), or TBS for 30 min at 25°C. Following treatment, PRP or washed platelets were diluted 1:20 in PBS containing either anti-human or anti-mouse FITC CD62p (Thermo Fischer Scientific, 1:67), depending on the species origin of the platelets. The samples were incubated for 30 min and subsequently analyzed using flow cytometry (Cytek Biosciences, Amnis CellStream).

### Murine passive ITP model

Mice were bled from the saphenous vein before the start of the experiment in EDTA Minivettes (Sarstedt AG & Co. KG) and platelets counts were measured using a Diatron Abacus - Model Vet5 (Diatron Budapest, Hungary.) Thrombocytopenia was induced in male BALB/c mice by intraperitoneal administration of rat anti-mouse CD41 (Becton Dickinson, clone MWReg30) at 3 µg per mouse. Subsequently, with no delay, mice were intraperitoneally injected with saline, 50 µM VLP, or 50 µM s4. Platelet counts were measured

24 h after the antibody injection as described above. At the end of the experiment, mice were sedated and exsanguinated by cardiac puncture into 100 µL citrate-phosphate-dextrose solution (Merck Life science). Blood was centrifuged for 2 x 10 min at 2500 x g, and plasma was frozen at -80°C for subsequent analysis. The body (rectal) temperature was measured before rat anti-mouse CD41 injection as well as 30 min and 24 h after. All mice were housed at the BMC Conventional Animal Facility at Lund University, Lund, Sweden. All animal studies were approved by the animal ethics committee of Lund University (Permit No: 5.8.18-00493/2022) in accordance with the guidelines of the Swedish National Board of Agriculture and the European Union directive for the protection of animals used in science.

### Production and purification of DNA origami flatsheet

A 3-tessellation DNA origami flatsheet was designed as previously described.^67^ The self-assembly of the structure was performed by mixing 10 nM scaffold DNA plasmid p8064 (Tilibit), 100 nM staple strands (Integrated DNA Technologies), 1 µM biotinylated staples (Integrated DNA Technologies), 1 µM digoxigenin (DIG) staples (Integrated DNA Technologies) in a folding buffer [5.0 mM Tris pH 8.5, 1.0 mM EDTA, and 13 mM MgCl_2_]. The reaction was run on a thermocycler (MJ Research PTC-225 Gradient Thermal Cycler) following a cycling protocol: step 1, 80.0 °C for 5 min; step 2, gradual cooling to 60.0 °C at 1.0 °C per minute; step 3 slow cooling to 20.0 °C at 0.5 °C per minute. Excess staples were removed by 100 kDa Amicon Ultra 0.5 ml centrifugal filter units (Merck). In brief, the sample were added to the filter units and washed 5x with 400 µl PBS supplemented with 13 mM MgCl2 by centrifugation for 2 min at 14,000 × g. The concentration of the purified DNA origami flatsheets was measured by a NanoDrop 2000 (Thermo Fisher Scientific).

### DNA-PAINT

#### Immunofixation of cells

Eighteen well glass bottom slides (Ibidi, µ-Slide) were coated 0.01 % poly-L-lysine (PLL) for 60 min at 37°C, washed 3x with distilled water, and dried for 30 min at 37°C. Serum starved 0.1 x 10^6^ THP-1 cells were treated with 0.5 μM scFc-VLPs, 0.5 μM scFc, 0.5 mg/ml HAIgG or 0.5 mg/ml IgG for 30 min at 37°C. Next, the cells were washed with PBS, resuspended in PBS and added to the PLL-coated glass slides for 5 min at 37°C. After surface adhesion, the cells were fixed with pre-warmed 4% PFA/PBS for 12 min at 25°C, washed 3x with PBS, blocked with blocking solution [3 % Bovine Serum Albumin (BSA)/0.1 % Triton X-100 in PBS] for 90 min at 25°C, and incubated with 5 μg/ml anti-human FcγRIIa (Origene, clone OTI9G5; 5 μg/ml) in blocking solution at 4°C overnight. Cells were then washed with PBS and incubated with FluoTag®-XM-QC anti-mouse IgG nanobody (Massive Photonics, 1:200 dilution in blocking buffer (Massive Photonics)) for 1 h at 25°C. After three washes with PBS, 80 nm gold nanoparticles (Sigma-Aldrich, 1:5 dilution in blocking buffer) were added to the cells for 10 min as drift markers. Cells were washed once with PBS and incubated with 5 nM ATTO 655-ssDNA (Massive Photonics) in an imager buffer (Massive Photonics). DNA-PAINT imaging was performed immediately after the addition of the imager strand.

#### DNA-PAINT imaging

DNA-PAINT imaging was performed using a Nikon ECLIPSE Ti-E microscope equipped with a Perfect Focus system (Nikon Instruments), iLAS2 circular total internal reflection fluorescence module (TIRF) (Gataca Systems) and an oil-immersion objective (Nikon Instruments, CFI Plan Apo TIRF ×100, NA 1.49, Oil). Samples were imaged in TIRF mode with an OBIS 561 nm LS 150 mW laser (Coherent) and custom iLas input beam expansion optics (Cairn) for reduced field super-resolution imaging. The light beam was passed through a filter cube (Chroma Technology, 89901), comprising an excitation quadband filter, a quadband dichroic filter, and a quadband emission filter (Chroma Technology, ZET405/488/561/640x, ZET405/488/561/640bs, and ZET405/488/561/640m). Emission filter (Chroma Technology, ET595/50m) was used to spectrally filter the fluorescence light that was captured on an EMCCD camera (iXon Ultra 888, Andor). The applied ×1.5 auxiliary Optovar magnification, resulting in an effective pixel size of 87 nm per pixel. The system was controlled by Micro-Manager software v. 1.4 and images were acquired at 10,000 frames, 10 MHz readout frame, 200 ms exposure.

#### Image processing and cluster analysis

Raw images were reconstructed by Picasso software (https://github.com/jungmannlab/picasso). The ‘Localization’ module was used to obtain the *x,y* localization coordinates of the single molecular events. The minimum threshold of the net gradient was set to 1000 and maximum likelihood estimation algorithm was applied with convergence criterion 0.001 and maximum number of iterations 1000. A custom-written code was used to filter localizations based on their precision error and Gaussian fitting. Next, drift correction was performed stepwise by the ‘Render’ module. In brief, a redundant cross-correlation algorithm was applied with a segmentation value of 1000, followed by global drift correction by gold nanoparticles used as fiducial markers. Localizations that persisted over multiple consecutive frames were removed using Link localizations function from the ‘Render’ module, with max distance equal to the localization precision error and a maximum transient dark frame of 15. One to three randomly assigned regions of interest (ROI) with 8.7 µm diameter were selected per cell by the ‘Render’ module and further analyzed.

A trained neural network model 87B1441^68^ with 1000 nearest neighbors as input was used for localization analysis, as previously described.^69^ A custom-written code using a mean frame filter and a standard deviation filter was applied to remove unspecific signals of the imaging strands. Statistical analysis was performed using a two-tailed non-parametric Mann-Whitney test.

A custom-written Python code was used to compute distances between neighboring FcγRII oligomers (≥3 receptors). The process involved a nested loop that systematically traverses each cluster, comparing Euclidean distances with other clusters to identify the minimal distance between the edges of two clusters. Subsequently, each cluster was associated with its closest neighbor, along with the respective distance. The average median distance was then computed from these distances across all clusters.

#### DNA origami calibration

Eighteen-well glass slides (µ-Slide, Ibidi) were cleaned with isopropanol and dried with N_2_. The slides were coated with 1 mg/ml Biotin-BSA (Thermo Fisher Scientific) in buffer A [10 mM Tris-HCl, 100 mM NaCl, 0.05 % Tween 20 at pH 8] for 5 min at 25°C and washed 3x with buffer A. Next, 0.5 mg/ml Neutravidin (Thermo Fisher Scientific) was added for 5 min at 25°C, washed 3x with buffer A and 3x with buffer B [5 mM Tris-HCl, 10 mM MgCl2, 1 mM EDTA, 0.0 5% Tween 20 at pH 8]. The biotinylated DIG labeled DNA origami flatsheets at concentration 300 pM were incubated for 5 min, followed by three washes with buffer B. Next, 60 nM of mouse Anti-Digoxigenin IgG1kappa (Sigma-Aldrich, clone 1.71.256) and 6 nM FluoTag®-XM-QC anti-mouse IgG nanobody (Massive Photonics) were added for 45 minutes at 25°C in blocking buffer (Massive Photonics). After three washes with buffer B, 5 nM ATTO655-ssDNA (Massive Photonics) in an imager buffer (Massive Photonics) was added. Imaging, fitting, filtering, and drift correction were performed as previously described. Single binding sites (n=150) on DNA origami flatsheets, displaying triangular arrangement, were manually selected by the ‘Render’ module. The average number of localizations per spot was calculated. Discrimination between monomers, dimers and receptor oligomers was determined using the following equation: number of receptors = localizations per cluster/localizations per spot.

## Acknowledgements

This work was supported by the European Research Council under the European Union’s Seventh Framework Program (617711, A.I.T.), the Swedish Research Council (2020-01856, A.I.T., 2021-01310, J.W.S), Avtal om Läkarutbildning och Forskning (ALF, J.W.S), the Knut and Alice Wallenberg Foundation (KAW 2017.0114, A.I.T.), and the ENDFLU consortium (874650, I.A.) from the European Union H2020 funding framework. Parts of this work were facilitated by the following core facilities: SciLifeLab Advanced Light Microscopy (ALM) Facility, Biomedicum Flow Cytometry Core Facility, and Biomedicum Imaging Core. All cryo-EM data was collected in the Karolinska Institutet’s 3D-EM facility. We acknowledge Protein Production Sweden (PPS) for providing facilities and experimental support for scFc synthesis, and we would like to thank Nicklas Bonander for assistance with gel-running and image preparation. PPS is funded by the Swedish Research Council as a national research infrastructure.

## Author contributions statement

E.P. designed, carried out experiments, and analyzed the data. G.K. designed, performed, analyzed and interpreted the DNA-PAINT experiments. J.R. and K.J. designed, performed, collected, and interpreted all in vitro platelet experiments and all animal experiments. S.W. performed, analyzed and interpreted FCS experiments. N.E.J.M. and I.A. performed, analyzed and interpreted molecular modeling experiments. I.A. supervised molecular modeling experiments. M.H. supported the design, analysis, and interpretation of Cryo-EM experiments. E.A. designed, performed, analyzed and interpreted the SPR experiments. J.W.S. provided financial resources and designed, analyzed, interpreted and supervised all platelet-related and mouse experiments. E.P. and A.I.T. conceived the study. A.I.T. directed the study. E.P. and A.I.T. wrote the manuscript and all authors reviewed and edited the manuscript.

## Data availability

The supporting data for this study can be obtained by contacting the corresponding authors. Cryo-EM maps are accessible from the Electron Microscopy Data Bank (EMDB) with the following accession codes: EMD-19437 for VLP and EMD-19438 for scFc-VLP.

## Extended Data Figures

**Extended Data Fig. 1:**
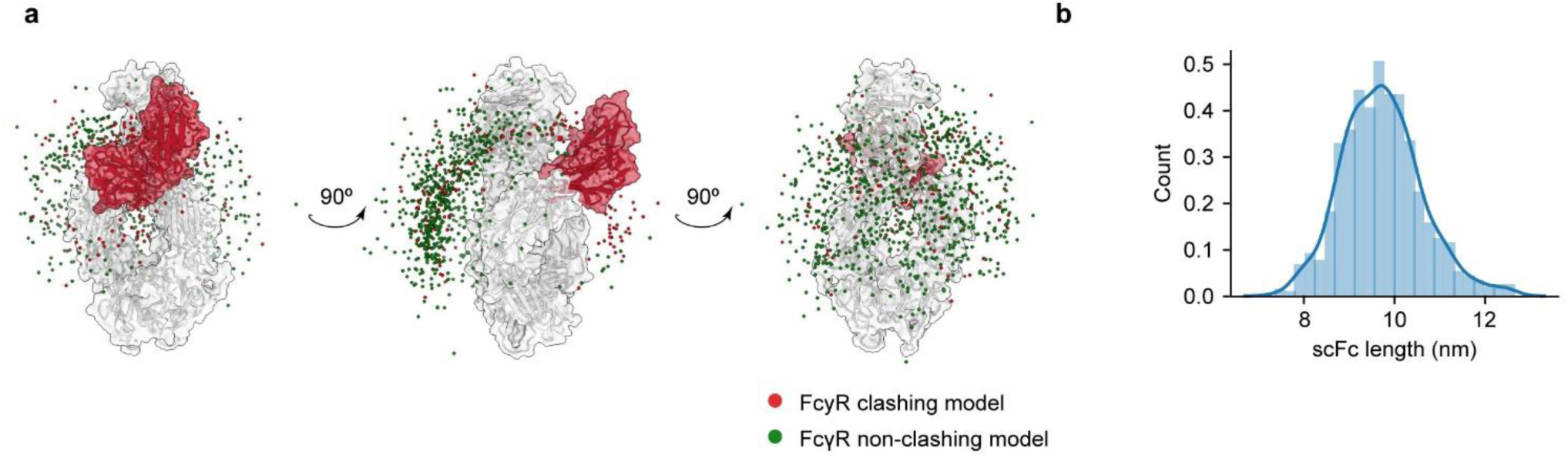
Computational characterization of scFc. **a,** Schematic of an ensemble of energetically plausible SNAPtag conformers linked to the Fc domain (depicted in a ghostly white) of the scFc. The center of mass (COM) of each conformer is denoted by a dot. Red dots highlight SNAPtag conformers clashing with the FcγRIIa (depicted in red), while green dots indicate non-interfering conformers. **b**, Distribution of scFc length. The scFc length is estimated based on the distance between the Fc and SNAPtag domains of the scFc, determined by the sum of their radii of gyration and the distance between their COMs. The mean scFc length is estimated to be 9.7 ± 0.9 nm.

**Extended Data Fig. 2:**
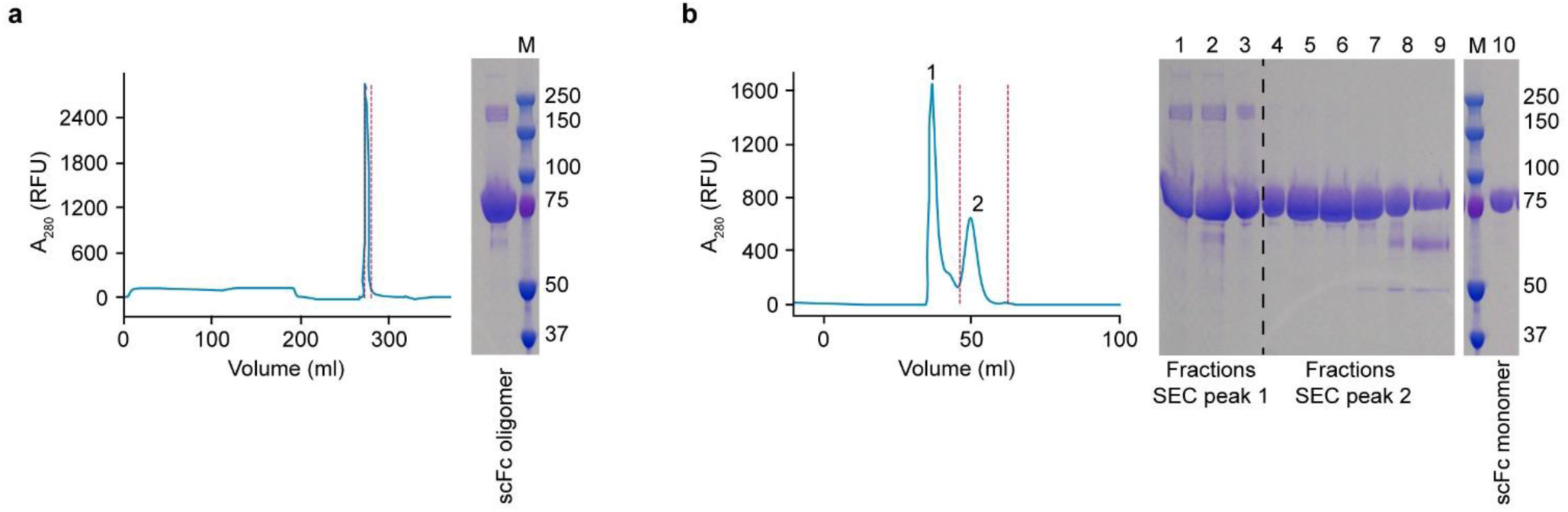
Purification of scFc. **a,** scFc was purified by affinity chromatography from cleared HEK 293F cell supernatant. The scFc retained its binding to FcRn and was eluted from Protein G affinity column. The red dotted lines in the plot enclose the collected elution fractions. Absorbance at 280 nm was monitored as a function of elution volume (ml) (left). Coomassie stained SDS-PAGE of scFc from collected elution fractions (right). **b,** SEC profile of scFc after Protein G affinity purification. Two distinct scFc forms were eluted: 1) scFc dimer, and 2) scFc monomer. Red dotted lines enclose the collected elution fractions (left). Coomassie stained SDS-PAGE of scFc dimer (L1-L3), scFc monomer (L4-L10) and the concentrated and filtered final scFc product (L10) (right). The molecular weight markers lane (M) is the same in **a** and in **b.**

**Extended Data Fig. 3:**
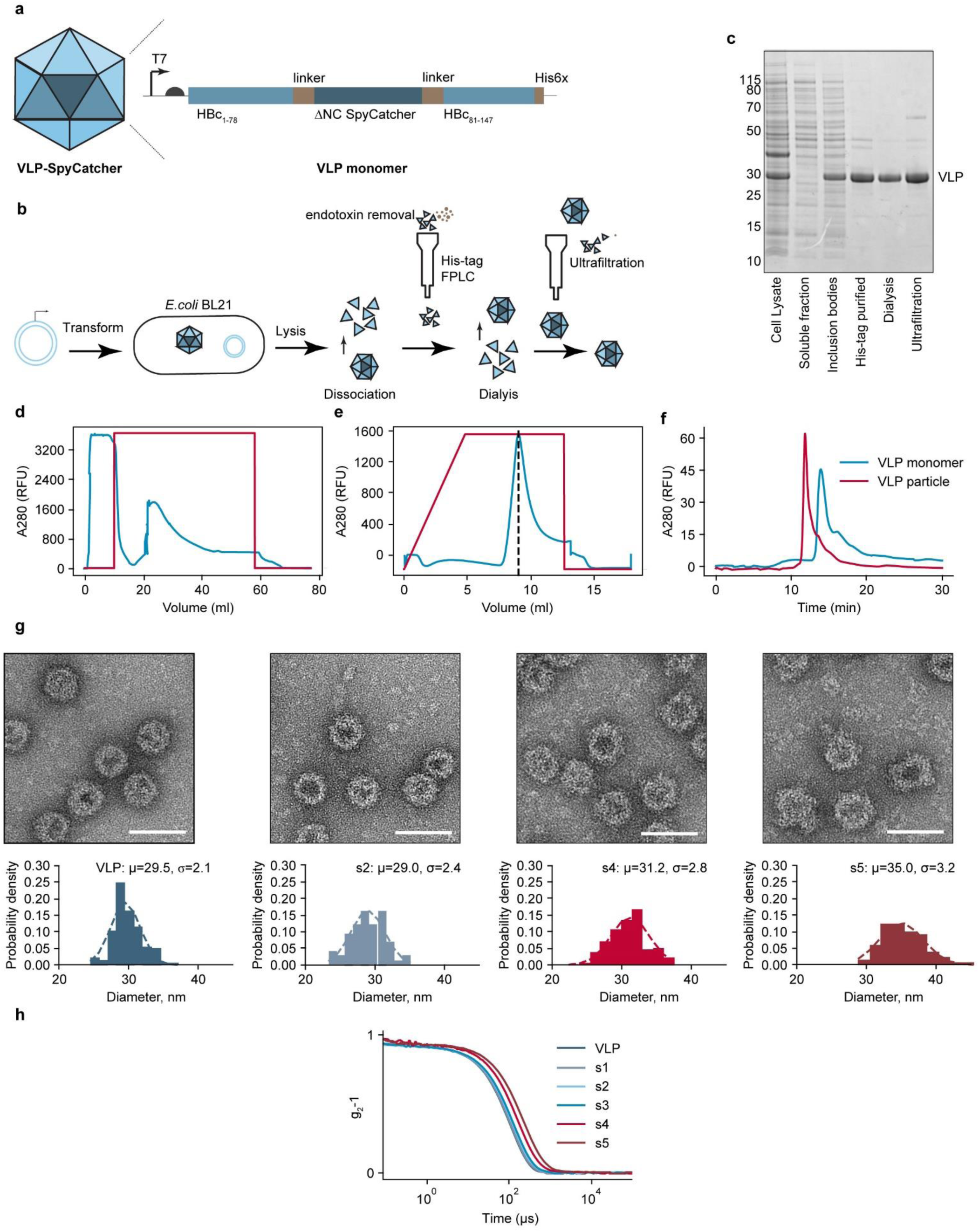
Design, production, and characterization of the VLP-SpyCatcher. **a,** Design of the VLP monomer. The VLP monomer, based on the Hepatitis B core (HBc) protein sequence, features a C-terminal truncation to prevent nonspecific nucleic acid binding to the VLP core and a ΔNC SpyCatcher, genetically fused by flexible flanking linkers within the HBc. VLP expression in *E. coli* BL21 is driven by the T7 promoter. **b,** Schematics illustrating key steps in the production and purification of VLP-SpyCatcher. **c,** Silver-stained denatured SDS-PAGE of total and soluble fractions, as well as the dissociated inclusion bodies fraction of SpyCatcher-VLP. Samples after affinity purification, dialysis, and ultracentrifugation (final product) were also resolved on the gel. **d** and **e,** Chromatograms of optimized affinity purification on ÄKTA FPLC for simultaneous VLP monomer purification and endotoxin removal. The red line depicts the protein absorption profile, and the blue line depicts the percentage of elution buffer. **d,** Binding and extensive washing with Triton X-114-containing binding buffer. **e,** Elution of the purified VLP monomer. **f,** SEC profile of dissociated VLP monomers and VLP sample after ultracentrifugation, showing distinct elution Times. **g**, Representative TEM images of VLP and scFc-VLPs (s*2*, s4, s5). The mean particle diameter is presented for each sample (n>50, mean ± std). Scale bars: 50 nm. **h**, DLS correlation functions measured at 25°C of VLP and scFc-VLPs (s1, s2, s3, s4, s5).

**Extended Data Fig. 4:**
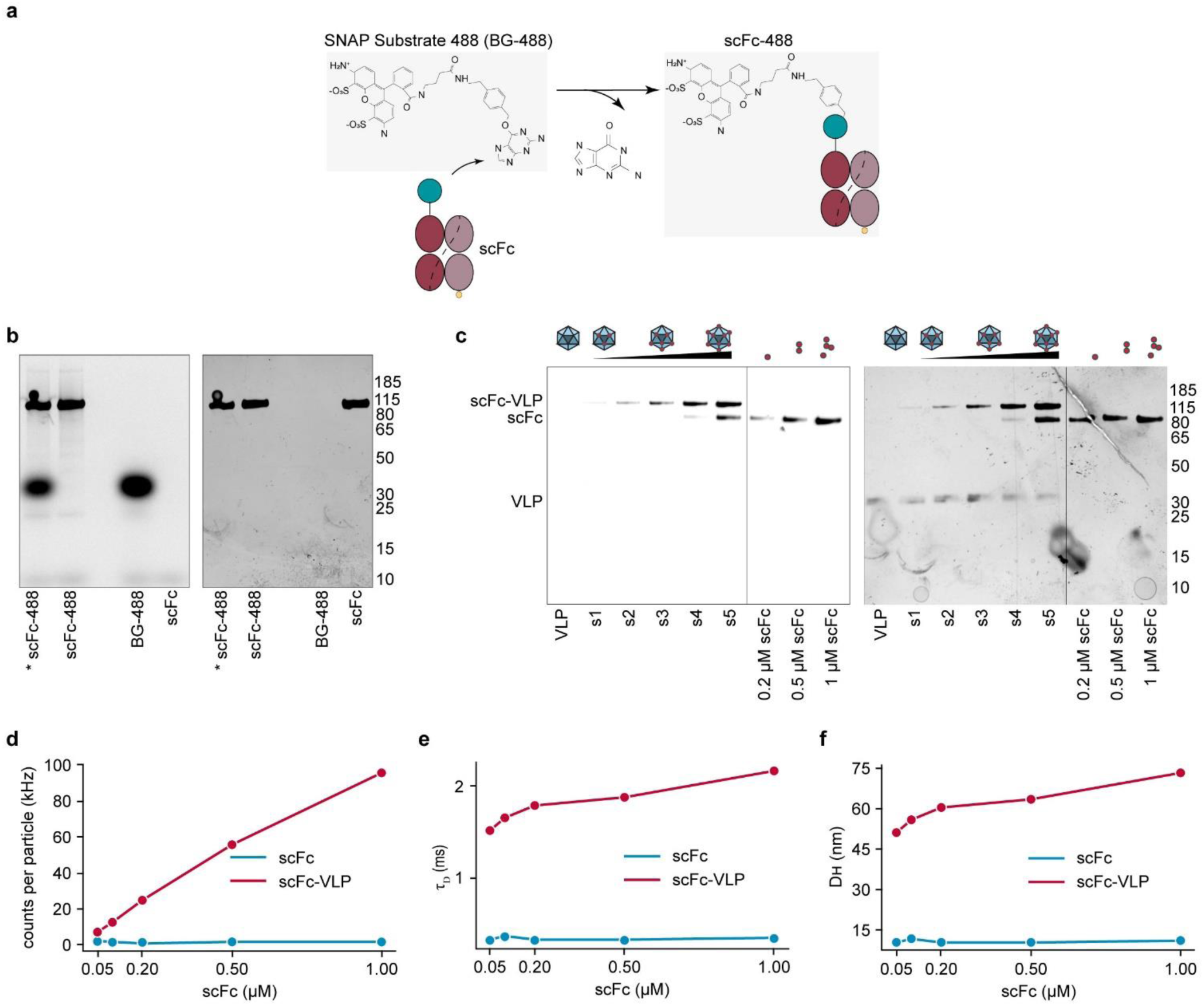
Synthesis and biophysical characterization of fluorescently labeled scFc and scFc-VLPs. **a**, Schematics of the site-specific conjugation between a scFc, containing a SNAPtag, and a SNAP Substrate 488, a BG-modified ATTO 488 dye (BG-488). **b,** Evaluation of the scFc-488 conjugation using SDS-PAGE, imaged using both fluorescent detection (left) and silver staining (right). * indicates unpurified scFc-488 containing an excess of free dye. **c,** SDS-PAGE analysis of VLP, scFc-VLPs (s1, s2, s3, s4, s5), and scFc at three different concentrations (0.2, 0.5, and 1 μM). Purified scFc-488 was used for scFc-VLP conjugation. Visualization was performed using fluorescent detection (left) and silver staining (right). **d-f**, FCS analysis of scFc-VLPs (red line) and scFc-488 (blue line) at different concentrations, corresponding to the scFc concentrations used for the assembly of the different scFc-VLP variants. **d,** Small variations in the molecular brightness (1.6 ± 0.2 kHz) were measured for all scFc concentrations and a steady increase in the molecular brightness with scFc valency was observed for the scFc-VLPs. **e,** The diffusion time (τ_D_) was constant for all scFc concentrations in the range of 0.3 ± 0.02 ms, while a steady increase was observed for the scFc-VLPs with increasing scFc concentration. **f,** The hydrodynamic diameter, which is proportional to the diffusion time, therefore also exhibited a steady increase in scFc-VLPs with increasing scFc concentration and was constant for the scFc concentrations.

**Extended Data Fig. 5:**
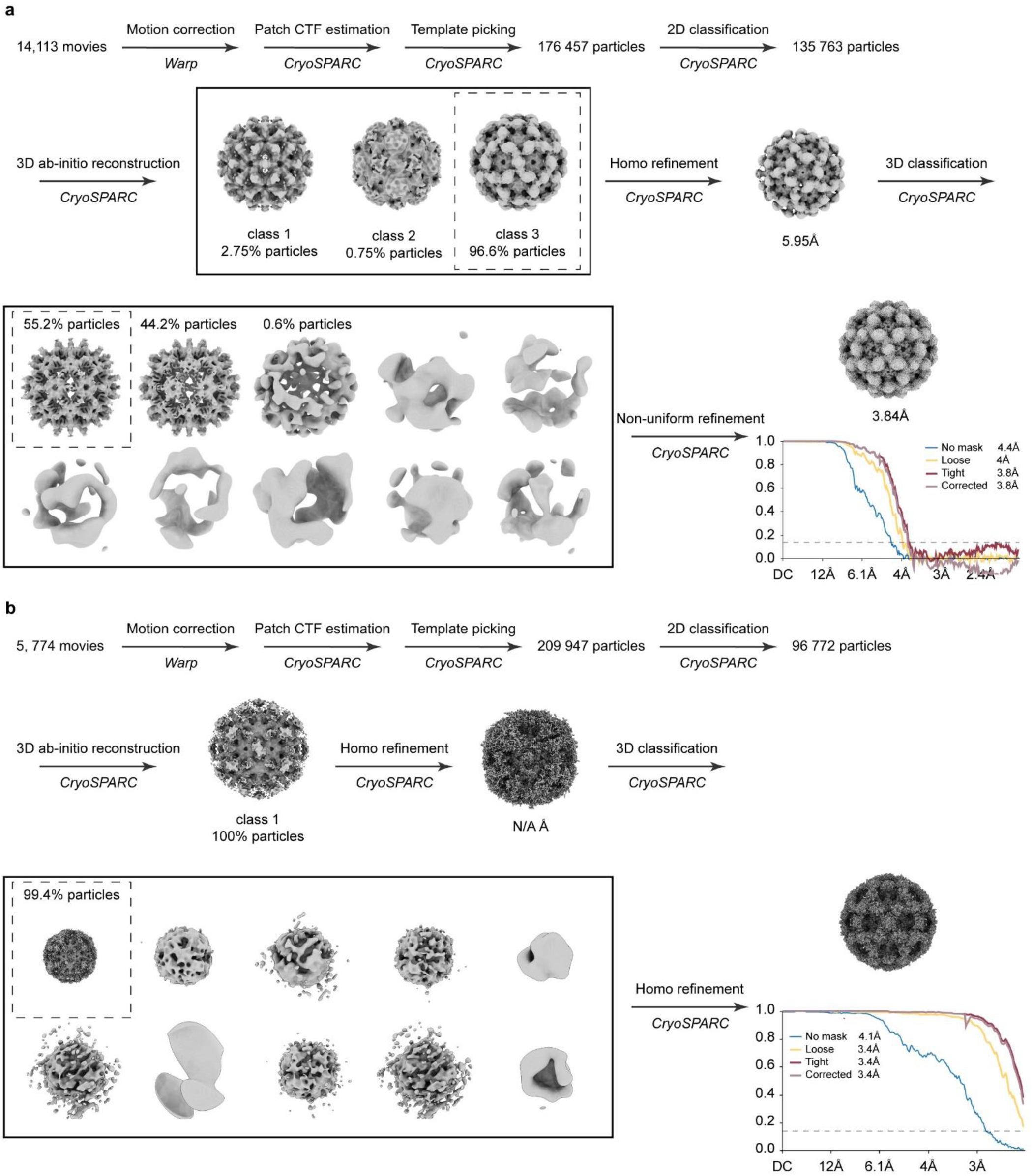
Cryo-EM data processing pipeline. **a** and **b,** Flow chart depicting data processing steps for icosahedral reconstructions of **(a)** VLP and **(b)** scFc-VLP. The Fourier Shell Correlation (FSC) value for the scFc-VLP does not reach the FCS gold standard, potentially due to limitations imposed by the Nyquist limit, given the low magnification employed during the image acquisition.

**Extended Data Fig. 6:**
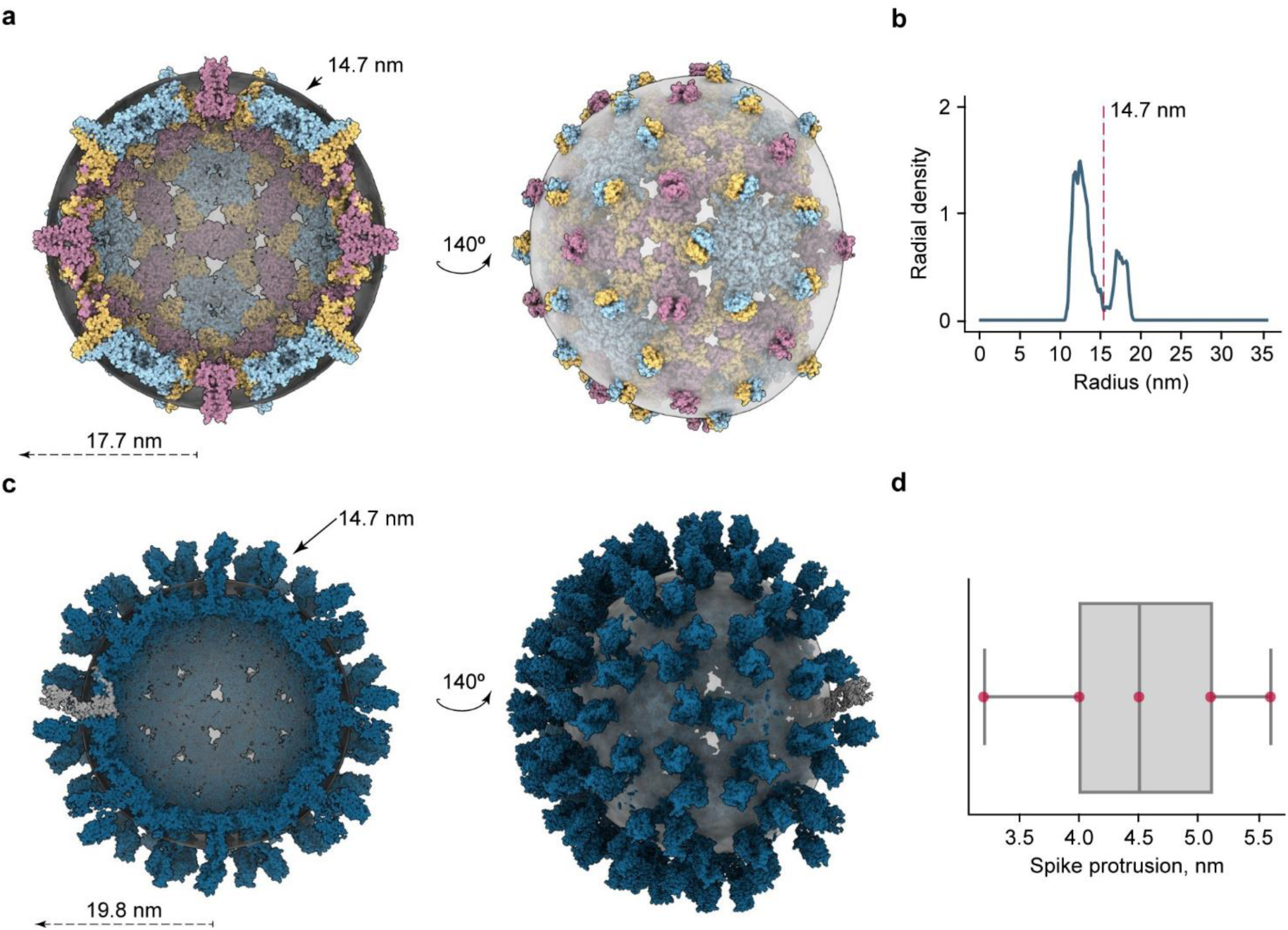
Structural characterization of VLP density models. **a**, Front and side view of the bottom half of HBV (PDB ID 6BVN) density map with color-annotated chains (depicted in blue, yellow, pink). **b,** Radial density profile of the HBV map, highlighting a local density minimum at 14.7 nm. **c,** Bottom half of the VLP structural model (VLP M1) featuring a single SpyCatcher-HBc monomer, depicted in gray. **d,** Boxplot of VLP spike protrusion distance. The distance is determined as the difference between the estimated outer radii of the VLP predictive models and the local HBV radial density minimum (n=5, median±std). A spherical surface (depicted in gray) with 14.7 nm radius, equal to the local HBV radial density minimum, is aligned and centered on both the HBV map **(a)** and the VLP M1 map **(d)**. The protruding spikes measure approximately 3 nm and 5.1 nm, respectively.

**Extended Data Fig. 7:**
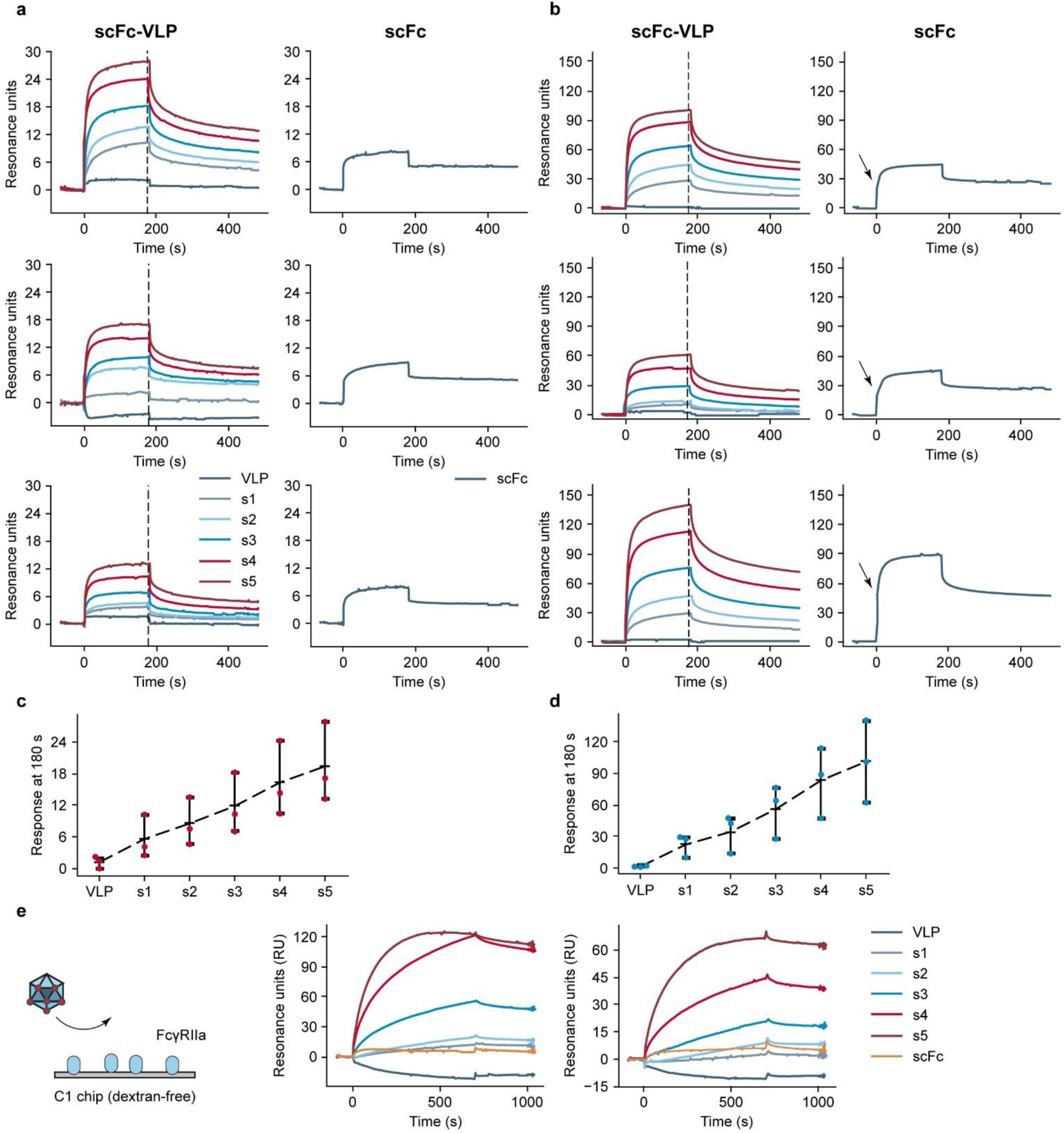
Profiling the binding of scFc and scFc-VLPs to FcγRIIa immobilized onto CM5 or C1 SPR surfaces. **a** and **b,** Binding traces of scFc-VLPs and scFc to CM5 chip immobilized with low levels (∼350 RU) **(a)** and high levels (∼1000 RU) **(b)** of ECD-FcγRIIa. Analyte concentration was 1.3 *µ*M. Each row represents an independent experiment. The scFc binding trace displayed an atypical behavior marked by the presence of an inflection point (annotated with a black arrow on the scFc sensorgrams), suggesting distinct interaction modes when the scFc binding was assessed on CM5 chips immobilized with high levels of FcγRIIa ECD. This behavior, distinct from that observed for scFc-VLPs, could be explained by the different accessibility of scFc to FcγRIIa immobilized on the dextran matrix compared with scFc-VLPs. **c** and **d,** Summary graphs of the scFc-VLPs response levels at 180 s (annotated as a dashed black line in the sensorgrams on **a** and **b**) for low (**c)** and high (**d)** immobilization levels of FcγRIIa. (n=3, mean ± std). **e,** Avidity effects exhibited by scFc-VLPs on dextran-free C1 chip. Binding of scFc, VLP, and scFc-VLPs (s1, s2, s3, s4, s5) to ECD-FcγRIIa immobilized on a dextran free C1 sensor chip, suitable to probe the binding of multivalent and high molecular weight complexes to immobilized proteins. The scFc binding was negligible and the binding trace had a similar shape compared with the other samples. Two independent experiments are shown.

**Extended Data Fig. 8:**
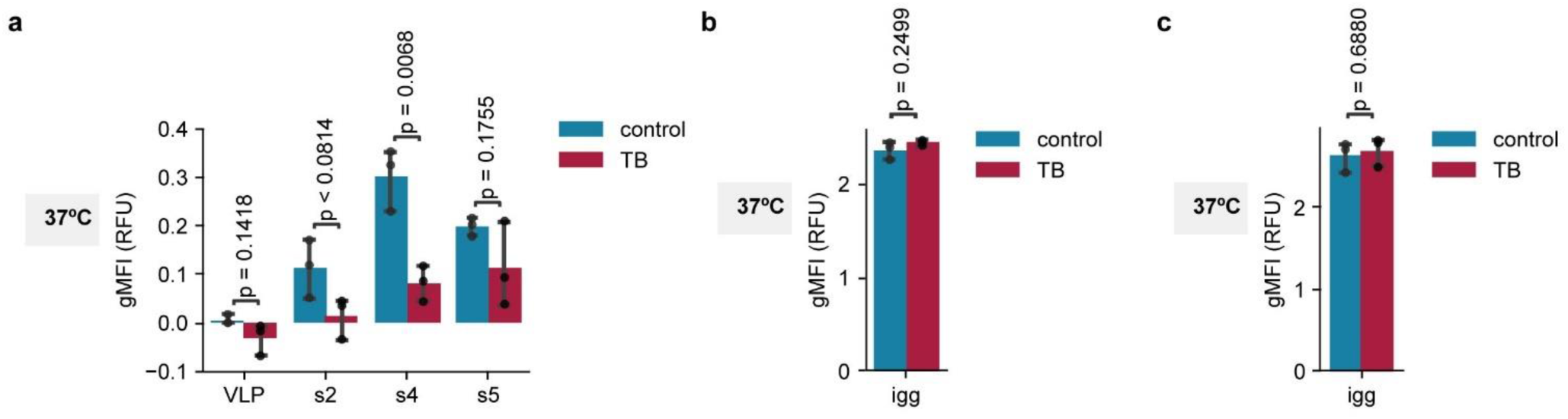
scFc-VLPs bind stably to the plasma membrane of THP-1 cells. **a,** Flow cytometry analysis of THP-1 cells treated with GFP-labeled VLPs or scFc-VLPs (s2, s4, s5) for 3 hours, washed with PBS, and left untreated (control) or incubated with TB. **b** and **c,** Flow cytometry analysis of THP-1 cells treated with IgG coated FITC-labelled beads for 30 min (**b)** or 3h (**c),** washed with PBS, and left untreated (control) or incubated with TB. The green fluorescent signal was measured shortly after the TB addition. Experiments were performed at 37°C. TB, Trypan Blue. P values were determined by one-way ANOVA followed by Tukey post-hoc test (n=3, median ± std).

**Extended Data Fig. 9:**
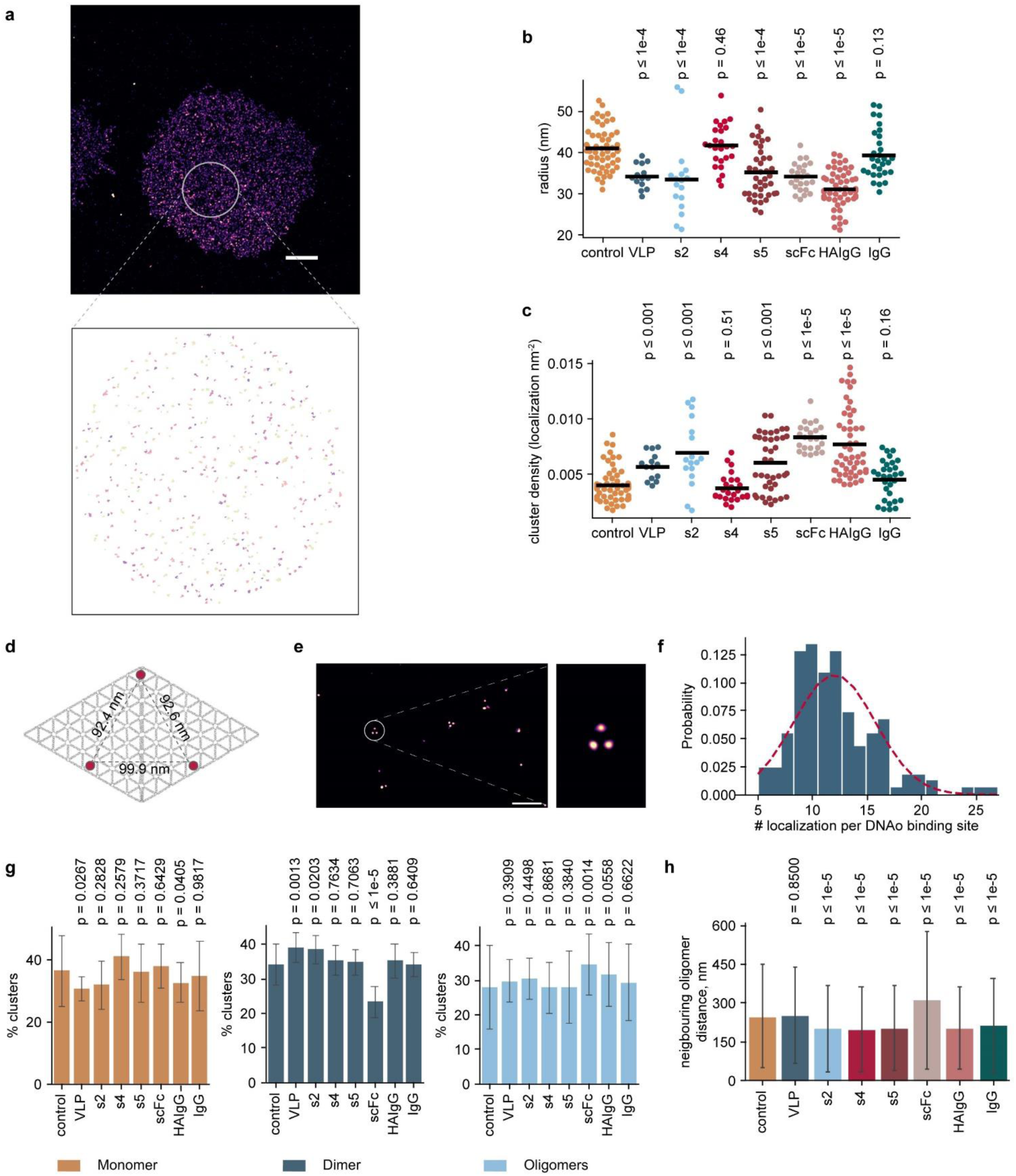
Analysis of FcγRII clustering in THP-1 cells. **a,** DNA-PAINT image of a THP-1 cell stained for FcγRII (upper panel). Scale bar: 5 µm. Higher magnification of a randomly selected region of interest (ROI) after image processing, showing distinct FcγRII localization clouds (lower panel). The diameter of the selected ROI is 8.7 mm. **b** and **c**, Characterization of FcγRII clusters in cells treated with PBS (control), VLP, scFc-VLPs (s2, s4, s5), scFc, HAIgG, or IgG. Plots of cluster radius (**b)** and cluster density (**c)**. P-values determined by a two-tailed Mann–Whitney test. **d**, DNA origami flatsheet with 3 designed binding sites (depicted as red spots). Distances between individual sites are annotated on the schematic. These structures were used to calculate the number of molecules per localization cluster, facilitating the quantification of FcγR numbers per cluster on THP-1 cells. **e**, DNA-PAINT image of the structure. The inset depicts a magnified view of the image showing an intact structure. Scale bar: 0.5 µm. **f**, Distribution of the number of localizations measured in one DNA origami (DNAo) binding site. A Gaussian fit to the data is depicted as a red line. The peak at 12.02 ± 3.74 corresponds to the average number of localizations per spot. This value was used to quantify the number of receptors in the detected FcγRII receptor clusters on THP-1 cells. **g,** Percentage of FcγRII clusters estimated to be monomers, dimers and oligomers (≥3 receptors) in THP-1cells left untreated (control) or treated with VLP, scFc-VLPs (s2, s4, s5), scFc, HAIgG, IC, or IgG for 30 min at 37°C. **h,** Estimation of the distances between the edges of neighboring FcγRII oligomers (> 3 receptors) on THP-1 cells treated with PBS (control) or with VLP, scFc-VLP (s2, s4, s5), scFc, HAIgG, and IgG. Data is presented as median, and error bars indicate 5-95 percentile. Statistical analysis was performed by two-tailed non-parametric Mann-Whitney test, P-values calculated based on the comparison between the control and the indicated group. All samples, except VLP, show significant differences compared with the control. No statistical significance was observed between the scFc-VLPs and HAIgG. Data is presented as median and error bars indicate 5-95 percentile. Statistical analysis was performed by two-tailed non-parametric Mann-Whitney test, P-values calculated based on the comparison between the control and the indicated group.

**Extended Data Fig. 10:**
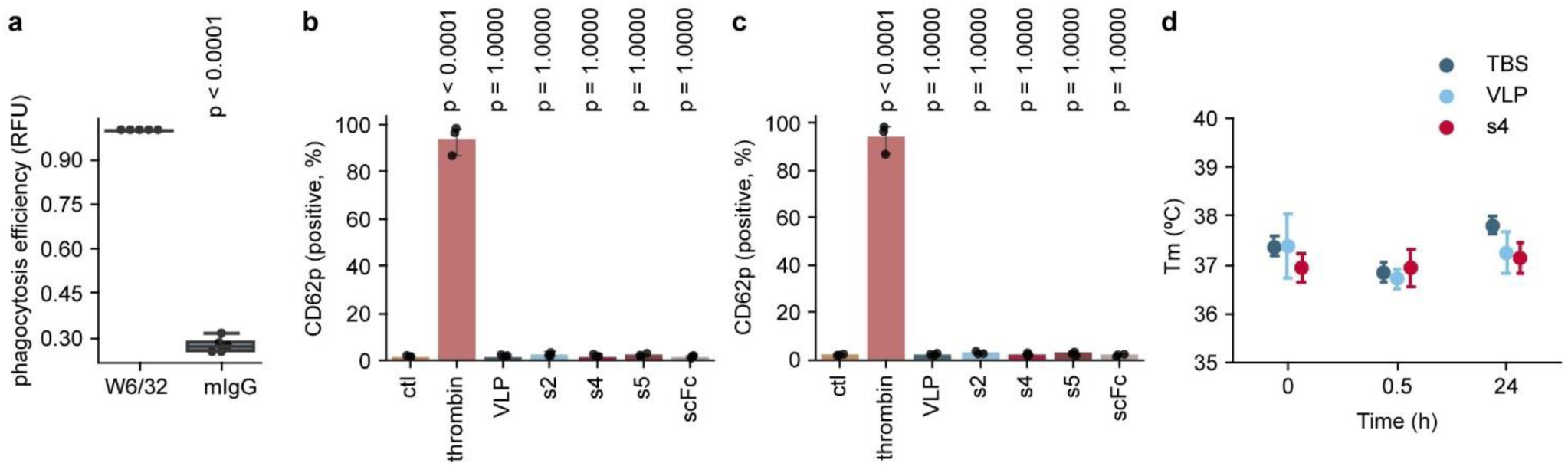
*Ex vivo* functional characterization and *in vivo* effects of scFc-VLPs. **a,** Flow cytometry depicting the phagocytosis efficiency of THP-1 cells incubated with platelets opsonized with anti-HLA (W6/32, positive control) or mouse IgG (mIgG, negative control). (n=5, mean ± std). **b** and **c**, Flow cytometry analysis of CD62p expression in mouse platelets left untreated (ctl) or treated with thrombin (positive control), VLP, scFc-VLPs (s2, s4, and s5), or scFc, in the absence **(b)** or presence **(c)** of mouse plasma. (n=3, mean ± std). **d**, Body temperature measurements in mice at 0, 0.5, and 24 h after i.p. injection of anti-CD41 antibody (n=5, median ± std). Statistical analysis was performed using one-way ANOVA and Tukey post-hoc test for **a-c**. The P-values are calculated based on comparison between the positive control (**a**) and the ctl (**b, c**).

