## Supplementary Figure 1 for "Engineering multivalent Fc display for FcγR blockade"

### Supplementary Information:

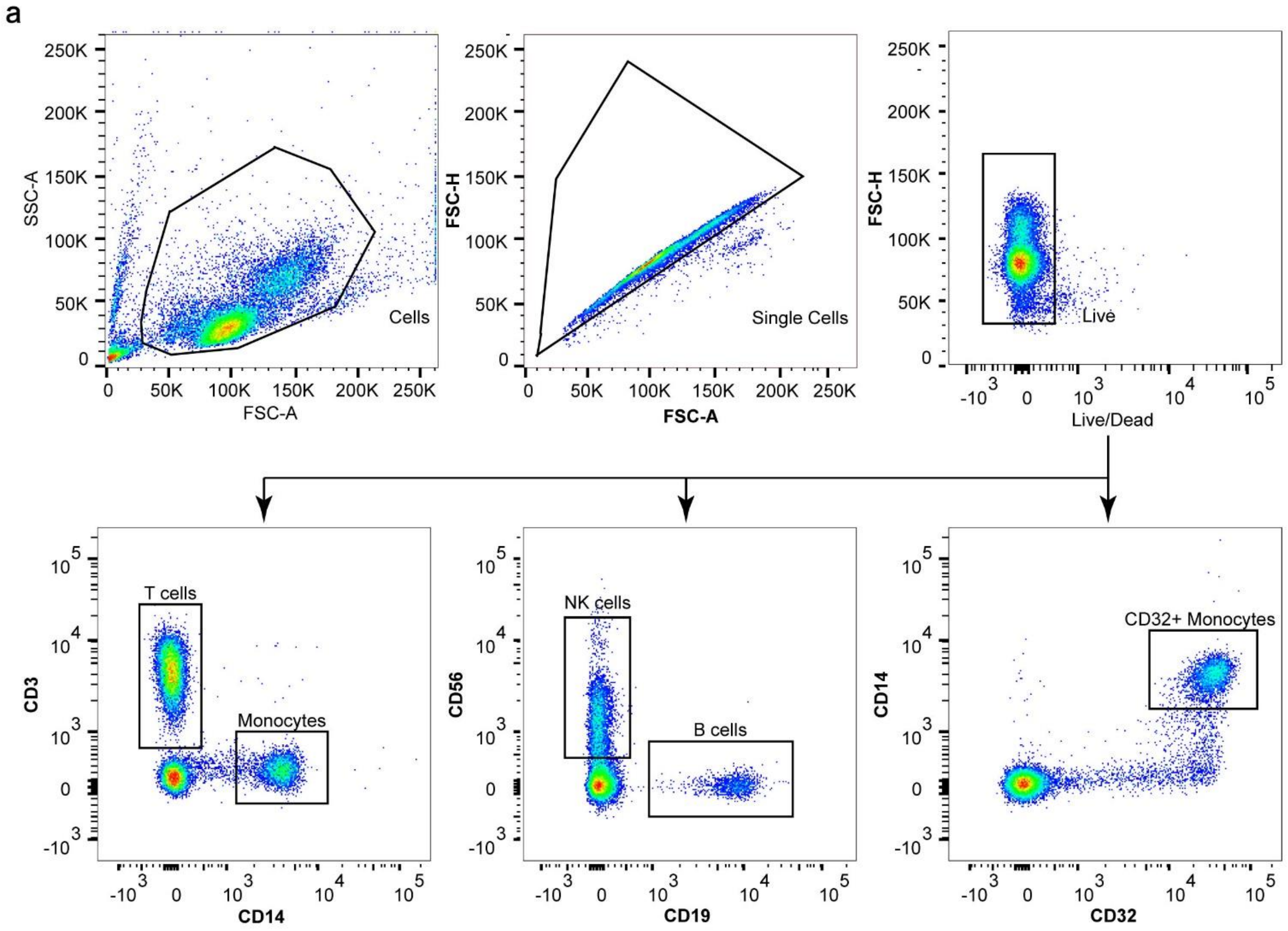


**Supplementary Fig.1: Gating strategy for phenotypic characterization of PBMCs**. Lymphocytes were identified based on their forward and side scatter area profiles (FSC-A and SSC-A, respectively) and single cells were gated based on their forward scatter height and area (FSC-H and FSC-A, respectively). Live cells were gated based on Live/Dead marker staining. Different cell populations were identified based on: 1) CD3 and CD14 labeling as T cells (CD3+) or monocytes (CD14+), 2) CD56 and CD19 labelling as B cells (CD19+) and NK cells (CD56+). We further gated the monocytes as CD32+CD14+ cells.
