## Supplementary Table 1 for "Engineering multivalent Fc display for FcγR blockade"

| | VLP | scFc-VLP |
|---|---|---|
| Microscope | FEI Krios G3i | |
| Voltage (kV) | 300 | |
| Detector | Bioquantum K3 | |
| Recording mode | Counting | |
| Magnification | 165,000 | 75,000 |
| Defocus range (μm) | 0.5-1.3 | 0.5-1.3 |
| Movie micrograph pixel size (Å) | 1.01 | 1.7 |
| Dose rate ($e^{-}$/Å$^2$/s) | 43 | 30 |
| Data collection automation software | EPU | |
| Image processing software | cryoSPARC v4.2.1 | |
| Number of frames per movie micrograph | 60 | 30 |

**Table S1 Cryo-EM settings**
